# Ketamine induces multiple individually distinct whole-brain functional connectivity signatures

**DOI:** 10.1101/2022.11.01.514692

**Authors:** Flora Moujaes, Jie Lisa Ji, Masih Rahmati, Joshua Burt, Charles H. Schleifer, Brendan Adkinson, Aleksandar Savič, Nicole Santamauro, Zailyn Tamayo, Caroline Diehl, Antonija Kolobaric, Morgan Flynn, Nathalie M. Rieser, Clara Fonteneau, Terry Camarro, Junqian Xu, Youngsun T. Cho, Grega Repovš, Sarah K. Fineberg, Peter Morgan, Erich Seifritz, Franz X. Vollenweider, John Krystal, John D. Murray, Katrin H. Preller, Alan Anticevic

**Author notes:** Both authors contributed equally to this work (Flora Moujaes, Jie Lisa Ji). Co-senior authors (Katrin H. Preller, Alan Anticevic). Correspondence &.

## Abstract

Ketamine has emerged as one of the most promising therapies for treatment-resistant depression. However, inter-individual variability in response to ketamine is still not well understood and it is unclear how ketamine’s molecular mechanisms connect to its neural and behavioral effects.

**Methods:** We conducted a double-blind placebo-controlled study in which 40 healthy participants received acute ketamine (initial bolus 0.23 mg/kg, continuous infusion 0.58 mg/kg/hour). We quantified resting-state functional connectivity via data-driven global brain connectivity, related it to individual ketamine-induced symptom variation, and compared it to cortical gene expression targets.

**Results:** We found that: i) both the neural and behavioral effects of acute ketamine are multi-dimensional, reflecting robust inter-individual variability; ii) ketamine’s data-driven principal neural gradient effect matched somatostatin (SST) and parvalbumin (PVALB) cortical gene expression patterns in humans, implicating the role of SST and PVALB interneurons in ketamine’s acute effects; and iii) behavioral data-driven individual symptom variation mapped onto distinct neural gradients of ketamine, which were resolvable at the single-subject level.

**Conclusions:** Collectively, these findings support the possibility for developing individually precise pharmacological biomarkers for treatment selection in psychiatry.

**Funding:** This study was supported by NIH grants DP5OD012109-01 (A.A.), 1U01MH121766 (A.A.), R01MH112746 (J.D.M.), 5R01MH112189 (A.A.), 5R01MH108590 (A.A.), NIAAA grant 2P50AA012870-11 (A.A.); NSF NeuroNex grant 2015276 (J.D.M.); Brain and Behavior Research Foundation Young Investigator Award (A.A.); SFARI Pilot Award (J.D.M., A.A.); Heffter Research Institute (Grant No. 1–190420); Swiss Neuromatrix Foundation (Grant No. 2016–0111m Grant No. 2015 – 010); Swiss National Science Foundation under the framework of Neuron Cofund (Grant No. 01EW1908), Usona Institute (2015 – 2056).

## Introduction

Over the last two decades, ketamine has emerged as one of the most promising therapies for treatment-resistant depression (TRD) ((1)). In March 2019 the Food and Drug Administration approved the use of intranasal esketamine for TRD in adults. Despite the wealth of research into ketamine’s molecular and systems-level neural effects ((2, 3)), two key knowledge gaps remain. First, inter-individual variability in the acute behavioral and neural response to ketamine has not yet been fully studied. Addressing this gap has direct clinical implications, as studies have shown that that a single ketamine infusion results in a response rate of around 65% in patients with TRD ((4)). Identifying markers of inter-individual variability in response to ketamine would allow us to predict the right treatment for each patient in line with a precision medicine approach. Second, it remains unclear how ketamine’s molecular mechanisms relate to neural systems-level and behavioral alterations. Bridging the gap between these multiple levels of analysis is paramount if we are to understand how pharmacological mechanisms map onto neural system-level predictive markers for personalized treatment selection in psychiatry.

There is evidence to suggest that ketamine results in robust inter-individual variability, as studies have shown individual differences in baseline molecular effects (e.g. NMDA receptor occupancy) and brain function predict the degree to which an individual experiences specific acute ketamine-induced symptoms ((5, 6)). Although ketamine’s neural systems-level functional alterations have been extensively characterised at the group level using both seed-based and whole-brain approaches, inter-individual variability in response to ketamine has been largely understudied. The majority of studies exploring ketamine’s systems-level alterations have used seed-based approaches ((7–19)). However, seed-based approaches cannot capture ketamine’s systems-level alterations in a data-driven way as they require the a priori selection of an area. Consequently, seed approaches may miss many of ketamine’s hypothesized system-level effects that alter connectivity across multiple areas ((20, 21)). Furthermore, while some studies have shown that ketamine results in altered thalamo-cortical and hippocampal-cortical activity, the direction of the effects are often inconsistent, which may in fact relate to individual variation ((10–12, 15, 19)). Put differently, inconsistent connectivity findings of ketamine’s acute effects could be a result of inter-individual variability in responses, which cannot be observed if studies focus only on the central tendency of the effect ((12, 15)). Indeed, Hack et al. (2021) found that ketamine resulted in dose-dependent individual variability in thalamic functional connectivity in healthy adults, with some individuals showing dose-dependent increases in functional connectivity while others showed dose-dependent decreases ((19)). However, this interim analysis is limited by its small sample size (N=7) and the use of a seed-based approach.

A limited number of studies have also characterised ketamine’s neural systems-level functional alterations using whole-brain approaches, demonstrating that ketamine results in robust brain-wide effects. For example, Driesen et al. (2013) used global brain connectivity (GBC) to show ketamine results in increased GBC in the cerebral cortex in healthy controls ((22)). However, this study lacked the spatial precision that is afforded by current state-of-the-art multi-band protocols (e.g. they found increases in GBC in large regions of white matter, which may be an artifact of large voxels and legacy volume-based alignment methods (22, 23)). Ketamine has also been shown to increase GBC in the pre-frontal cortex (PFC) in healthy controls specifically, and normalize the reduced GBC in the PFC in patients with major depressive disorder 24 hours post-ketamine ((24–26)). However, this finding failed to replicate in patients with depression 48 hours post-ketamine ((27)), suggesting a complex relationship between ketamine’s acute and delayed effects. In addition to GBC, graph theoretical approaches have demonstrated that ketamine induces a shift from a cortical to a subcortically-centred brain state, particularly the basal ganglia and cerebellum ((28)). Meanwhile, a nodal predictive model found that ketamine resulted in reduced connectivity within the primary cortices and the executive network, but increased connectivity between the executive network and the rest of brain ((29)). Finally, dynamic resting-state functional connectivity analyses showed that ketamine decreased connectivity both within the left visual network and inter-hemispherically between the visual networks ((30)). Together, these studies leave open the question of whether there may be robust inter-individual variability in ketamine’s neural systems-level, which has yet to be studied using a whole-brain connectivity data-driven approaches.

In order to address this gap, we studied inter-individual variability in response to acute ketamine administration in 40 healthy controls using a whole-brain data-driven multi-dimensional approach. We specifically tested the hypothesis that ketamine may affect individuals differently and thus result in multiple neuro-behavioral axes of acute effects. Put differently, we hypothesize ketamine will result in multiple significant individual dimensions of neuro-behavioral change, reflecting inter-individual variability in acute ketamine response that cannot be fully captured by the mean group-level approach. In addition, we will investigate the extent to which multi-dimensionality is a property specific to ketamine by directly comparing ketamine to two other psychoactive substances that have also shown preliminary clinical efficacy for the treatment of depression, but have very different pharmacological profiles: i) psilocybin, a preferential 5-HT2A and 5-HT1A agonist; and ii) LSD, which stimulates a wide range of serotonin and dopamine receptors ((31–36)).

In turn, we addressed another key knowledge gap concerning ketamine’s molecular mechanisms. Two mechanistic hypotheses have been proposed for ketamine’s cellular-level mechanisms of action: the direct and the indirect hypotheses ((37)). The direct hypothesis posits that ketamine directly antagonises N-methyl-D-aspartate (NMDA) receptors on glutamatergic neurons, causing the activity-dependent synaptic plasticity that underlies ketamine’s rapid antidepressant effects ((37)). Meanwhile, the indirect hypothesis proposes that ketamine first inhibits tonic-firing GABAergic interneurons via NMDAR blockade, which in turn leads to a burst of glutamate that drives synaptic plasticity ((37)). The majority of interneurons in the neocortex express one of three non-overlapping genetic markers: i) somatostatin (SST); ii) parvalbumin (PVALB); and iii) the ionotropic serotonin receptor 5HT3a ((38)). SST and PVALB interneurons have been shown to be particularly sensitive to ketamine as a result of their NMDA receptor subunit configuration ((39)). For example, SST and PVALB interneurons mainly express GluN2C (PVALB) and GluN2D (PVALB and SST) NMDAR subunits, which are only weakly blocked by magnesium ions at resting-membrane potential and so are more sensitive to ketamine ((40–42)). In contrast, glutamatergic pyramidal cells tend to express GluN2B NMDAR subunits, which are strongly blocked by magnesium ions at resting-membrane potential and so are less sensitive to ketamine ((43)). Furthermore, studies of patients with depression have reported reduction in both SST or PVALB interneurons in areas such as the hippocampus and the prefrontal cortex ((44–47)).

Here we build on previous research showing that specific cortical regions express different ratios of SST and PVALB interneurons ((48)). This is reflected in genome-wide atlases of spatially distributed gene expression profiles such as the Allen Human Brain Atlas (AHBA), in which SST shows higher expression levels in association relative to sensory regions, while PVALB shows the reverse pattern ((49)). We aim to connect ketamine’s hypothesized molecular mechanisms to its systems-level functional alterations by correlating ketamine-induced brain-wide changes in GBC with cortical SST/PVALB gene expression maps ((50)). Such an approach has already been used to highlight the critical role of 5-HT2A in LSD’s brain-wide connectivity effects ((51)). Overall, we hypothesize that: i) there will be an association between ketamine-induced changes in GBC and SST/PVALB gene expression profiles; and ii) that analysing ketamine’s neural effects using a multi-dimensional rather than a mean-based approach will enable us to more strongly track SST/PVALB gene expression profiles.

In addition to linking ketamine’s downstream molecular mechanisms to its neural systems-level functional alterations, it is also important we relate these changes to ketamine’s acute behavioral effects. For example, understanding the relationship between neural and behavioral changes could help inform predictions about how specific individuals will respond to ketamine in a clinical setting. At subanesthetic or antidepressant doses, ketamine produces transient changes in behavior, perception, and cognition that are comparable to the positive, negative, and cognitive symptoms seen in patients with psychosis-spectrum illness ((2)). Therefore, ketamine’s acute behavioral effects are typically captured using psychosis-related scales such as the Positive and Negative Syndrome Scale (PANSS), as well as cognitive tasks ((3)). Critically, however, investigations linking ketamine’s neural and behavioral alterations have failed to account for inter-individual variability. In order to address this gap, we will leverage an approach from the psychosis literature that involves using a data-driven dimensionality reduction method to quantify orthogonal dimensions that capture maximal individual variation of ketamine’s acute behavioral effect. In turn, we regress the resulting data-driven behavioral dimensions of ketamine’s acute effect onto its neural effects ((52)). We hypothesize that the principal axes of ketamine’s behavioral effects will: i) capture novel patterns of neural variation; and ii) track SST and PVALB neural gene expression patterns. In addition, we will also compare the data-reduced behavioral dimensions to established psychosis literature sub-scales (e.g. 3-factor and 5-factor PANSS models), and explore the extent to which this framework for mapping across multiple levels of analysis is resolvable at the single subject level.

Collectively, this study addresses major knowledge gaps in our understanding of ketamine’s neurobiology, by demonstrating: i) the neural and behavioral effects of ketamine are multi-dimensional, reflecting robust inter-individual variability; ii) ketamine’s data-driven principal neural gradient tracks SST and PVALB cortical gene expression patterns; and iii) ketamine’s data-driven principal behavioral gradient captures novel neural variation and provides an anchor point for both the neural systems-level effects and molecular mechanisms that is resolvable at the single subject level.

## Results

### Multidimensional neural effect of acute ketamine administration

In order to explore whether the neural effects of acute ketamine are uni-dimensional or multi-dimensional, we conducted a data-driven reduction procedure using a principal component analysis (PCA). The PCA was computed as follows: (1) in order to increase the signal to noise ratio, the neural data was first parcellated into 718 functionally-defined parcels (**Fig. S1**); (2) global brain connectivity (GBC) was computed across the parcels; (3) a Δ GBC map was created for each participant by calculating the difference between their GBC maps following ketamine and placebo infusion; and (4) a PCA was performed on the Δ GBC neural features (718 whole-brain parcels) across all subjects (N=40) (i.e. the input consisted of a 718 × 40 matrix). For comparison, a PCA was also performed separately on the placebo GBC neural feature and ketamine GBC neural features (**Fig. S2**). The PCA performed on the Δ GBC data revealed that ketamine produces five axes of neural alteration, as we identified five significant PCs (**Fig. 1A**) that together capture 42.1% of the total variance. However, as the first two significant PCs each capture a much greater proportion of the variance (> 10%) compared to each of the remaining three significant PCs (variance ~ 5%), we will focus on the first two PCs in the main text. Results for all five significant PCs are reported in the supplement (**Fig. S3, Fig. S4, Fig. S5**).

**Fig. 1.**
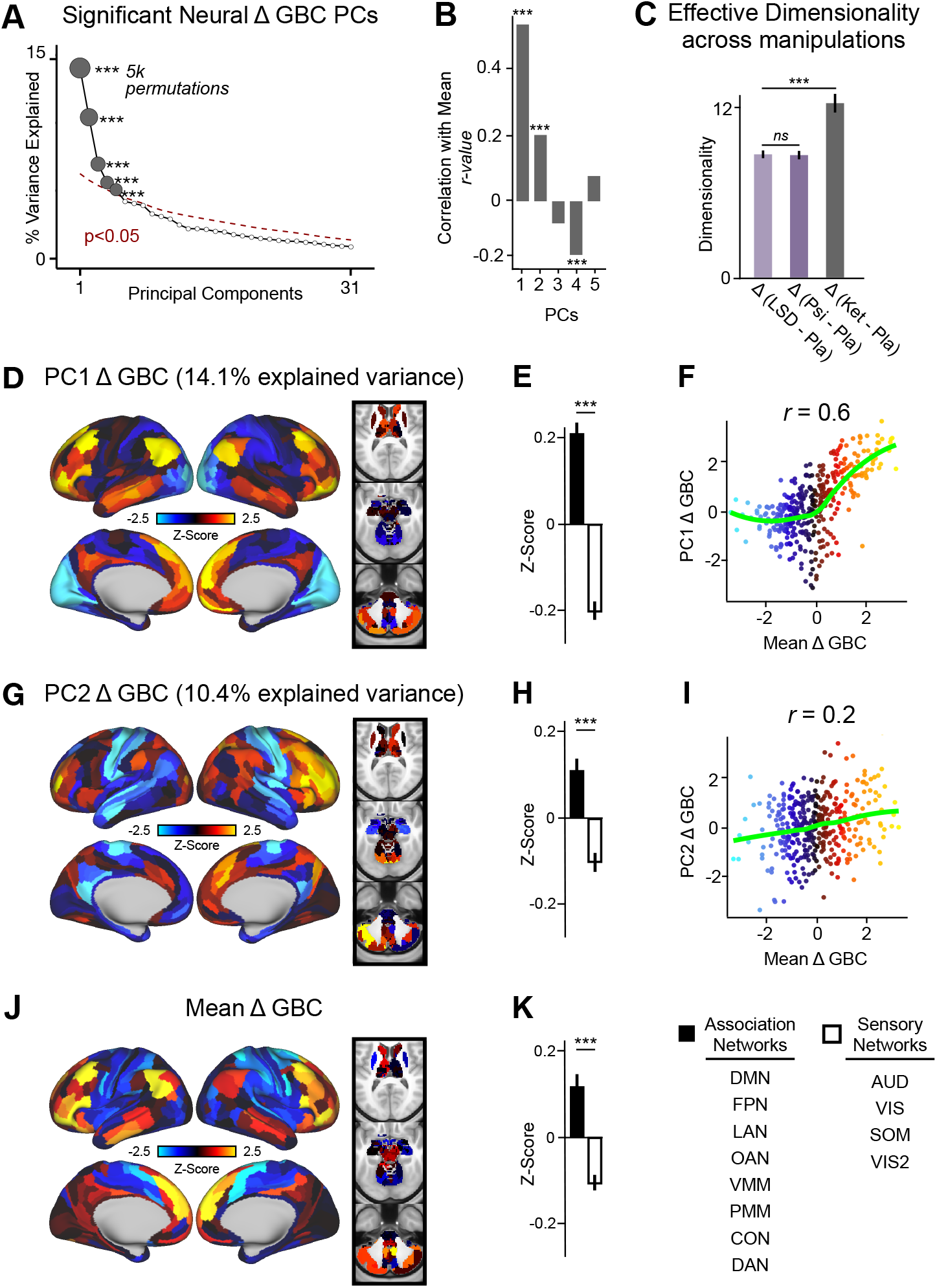
Multidimensional neural effect of acute ketamine administration. **(A)** Results of PCA performed on Δ GBC neural features (718 whole-brain parcel GBC) across all subjects (N=40). Screeplot showing the % variance explained by the first 31 (out of 39) Δ GBC PCs. The first 5 Δ GBC PCs (dark grey) were determined to be significant using a permutation test (p<.05, 5000 permutations). The size of each dark grey point is proportionate to the variance explained. Together, these 5 Δ GBC PCs capture 42.1% of the total variance in neural GBC in the sample. **(B)** Bar plot showing the correlation between each of the five significant Δ GBC PCs maps and the mean Δ GBC map. r values for PC1, PC2, and PC4 Δ GBC are significant (*** = p<.001, Bonferroni corrected). **(C)** Bar plot showing the effective dimensionality across three pharmacological conditions: Δ LSD (LSD - placebo), Δ psilocybin (psilocybin - placebo), and Δ ketamine (ketamine - placebo). (light purple = LSD, dark purple = psilocybin, dark grey = ketamine). There was a statistically significant difference between groups as determined by a one-way ANOVA (F(2,60)=564, p<.001). Post hoc tests revealed that dimensionality was significantly lower in the LSD (8.7 +/− 0.3, p<.001) and the psilocybin (8.6 +/− 0.3, p<.001) conditions compared to the ketamine (12.8 +/− 0.7) condition. There was no statistically significant difference in dimensionality between LSD and psilocybin (p=0.6). Error bars = standard deviation. **(D)** Unthresholded PC1 Δ GBC Z-score map (Z-scores computed across 718 parcels). PC1 Δ GBC explains 14.1% of all variance. Red/orange areas indicate parcels that have a high positive loading score onto PC1, while blue areas indicate parcels that have a high negative loading score onto PC1. **(E)** Bar plot showing the PC1 (Δ GBC Z-score across association (black) and sensory (white) network parcels. See bottom right for association and sensory network groupings. **(F)** Scatterplot showing the relationship across parcels between mean Δ GBC and PC1 Δ GBC maps (r=0.56, p<.001, n=718). **(G)** Unthresholded PC2 Δ GBC Z-score map (Z-scores computed across 718 parcels). PC2 Δ GBC explains 10.4% of all variance. Red/orange areas indicate parcels that have a high positive loading score onto PC2, while blue areas indicate parcels that have a high negative loading score onto PC2. **(H)** Bar plot showing the PC2 (Δ GBC Z score across association (black) and sensory (white) network parcels. **(I)** Scatterplot showing the relationship across parcels between mean Δ GBC and PC2 Δ GBC maps (r=0.23, p<.001, n=718). Green line indicates neither the positive or negative values are highly correlated. **(J)** Unthresholded mean Δ (ketamine - placebo) GBC Z-score map (Z-scores computed across 718 parcels). Red/orange areas indicate regions where participants exhibited stronger GBC in the ketamine condition, whereas blue areas indicate regions where participants exhibited reduced GBC in the ketamine condition, compared with the placebo condition. **(K)** Bar plot showing the mean Δ GBC Z-score across association (black) and sensory (white) network parcels.

In order to explore whether there is network specificity in the multi-dimensional ketamine effect, we first grouped parcels according to whether they belonged to sensory networks (i.e. primary visual, secondary visual, somatomotor, auditory) or association networks (i.e. default mode network, frontoparietal, language, orbito-affective, dorsal attention, ventro multimodal, posterior multimodal, cingulo-opercular). We then plotted the mean Z-score (standardized across the whole brain) across parcels belonging to association networks and parcels belonging to sensory networks for PC1 and PC2 Δ GBC. We found that PC1 and PC2 Δ GBC successfully differentiate between association and sensory networks (**Fig. 1D, E, G & H**). As PC1 and PC2 Δ GBC are bi-directional axes of variation, some individuals may exhibit increased GBC in association cortices and decreased GBC in sensory cortices, while others may show the opposite pattern (**Fig. S3A - D**). PC1 Δ GBC indexes variation most prominently in the secondary visual, default mode, and frontoparietal networks (**Fig. S4C**). In contrast, PC2 Δ GBC indexes variation most prominently in the somatomotor, frontoparietal, and dorsal attention networks and the hippocampus (**Fig. S4F**).

To test the hypothesis that these dimensions capture neural variance better than the mean effect, we also computed the mean Δ GBC map across participants (**Fig. 1J**). Though PC1 Δ GBC is moderately correlated with the mean Δ GBC (r=0.56), the subsequent significant PCs 2-5 are only weakly correlated with the mean Δ GBC (**Fig. 1B**). While this may seem self-evident given that a PCA enforces orthogonal axes, this point highlights that there is variation within the neural response to ketamine that is not sufficiently captured using a uni-dimensional approach. This is also evident when comparing the amount of variation captured by PC1 Δ GBC (*σ*^2^= 0.12) and the mean Δ GBC (*σ*^2^ = 0.05) (**Fig. S3B & L**). In addition, we found that PC1 Δ GBC and the mean Δ GBC maps do not have a linear relationship, as in **Fig. 1F** the sigmoid demonstrates that while the positive values are highly correlated, the negative values do not correlate.

We then ran a control analysis to explore whether this multi-dimensional neural effect is specific to ketamine or a property common to pharmacological neuroimaging in general. We compared the acute effect of ketamine administration to two drugs that also have anti-depressive effects, but very different pharmacological profiles: (1) psilocybin, a preferential 5-HT2A and 5-HT1A agonist; and (2) LSD, which stimulates a wide range of serotonin and dopamine receptors (33–36). For the LSD and psilocybin analyses we used the Δ GBC parcellated neural maps from two independent pharmacological neuroimaging datasets (51, 53). We calculated effective dimensionality for each drug, using a re-sampling method to ensure that sample size (which differed between the three datasets) was kept constant and did not bias the results (see Methods) (54–56). This revealed that ketamine was significantly higher in dimensionality than LSD and psilocybin (**Fig. 1C**, p<.001). In contrast, there was no difference in dimensionality between LSD and psilocybin (**Fig. 1C**, p=.6). We also ran a PCA on the psilocybin and LSD Δ GBC maps, which showed that both LSD and psilocybin result in fewer dimensions than ketamine (**Fig. S6**). Overall, this suggests that ketamine has a multi-dimensional effect on connectivity patterns in the brain, and that this effect shows some specificity to ketamine compared to other psychoactive substances such as LSD and psilocybin.

### PC1 Δ GBC tracks SST and PVALB neural gene expression topography, while mean Δ GBC does not

We hypothesized that analysing ketamine’s neural effects using a multi-dimensional approach rather than a mean-based approach will enable us to more strongly track SST/PVALB gene expression profiles. To test this, we evaluated the relationship between PC1-5 Δ GBC and the gene expression maps by first simulating 100,000 surrogate maps for each PC that preserved the PC Δ GBC map’s spatial auto-correlation properties. Secondly, for each PC we correlated both the SST and PVALB gene expression maps with each of the 100,000 surrogate maps to create a permuted distribution of r-values for SST and PVALB. Finally, for each PC we used the permuted distribution of r-values for SST and PVALB to estimate the probability of the observed correlation between the PC Δ GBC map and each gene expression map occurring by chance (**Fig. 2A**). For a more detailed outline of the gene expression analysis see **Fig. S7**. We found that as hypothesized, the multi-dimensional approach more strongly tracked SST/PVALB gene expression profiles, as PC1 and PC3-5 Δ GBC maps successfully captured the association between ketamine-induced changes in GBC and the cortical topography of SST and PVALB receptor gene expression while the mean Δ GBC map did not (**Fig. 2E & H, Fig. S5J, M & P**). For example, PC1 Δ GBC showed a significant positive correlation with the SST gene expression map (r=0.47, p<.001, 100,000 permutations) and a significant negative correlation with PVALB gene expression map (r=−0.22, p=.025, 100,000 permutations) well beyond what is expected by chance alone (**Fig. 2F**). In contrast, the the mean Δ GBC was not able to capture this association, as neither the SST or PVALB gene expression maps significantly correlated with the mean Δ GBC (**Fig. 2H**). Finally, to examine whether this effect holds for individual subjects, we selected the highest negative (subject 32) and highest positive (subject 28) loader onto PC1 Δ GBC and examined whether their individual Δ GBC maps also track SST and PVALB neural gene expression topography (**Fig. S8B**). We found a significant positive correlation between the highest positive loader’s Δ GBC map and the SST gene expression map (r=0.23, p=.009, 100,000 permutations), and no relation with the PVALB gene expression map (r=−0.07, p=.19, 100,000 permutations) (**Fig. S8D & E**). In contrast, we found a significant negative correlation between the highest negative loader’s Δ GBC map and the SST gene expression map (r=−0.22, p=.046, 100,000 permutations), and a borderline significant positive correlation with the PVALB gene expression map (r=0.16, p=.056, 100,000 permutations) (**Fig. S8G & H**). Overall, this demonstrates that using a multi-dimensional analytic approach is better than a mean approach at capturing the association between ketamine and interneuron markers SST and PVALB, which are hypothesized markers of mechanism. In addition, these results indicate that it is possible to derive the data-driven neural topography of a pharmacological effect, quantitatively relate it to gene expression, and evaluate this relationship using a non-parametric statistical permuted approach.

**Fig. 2.**
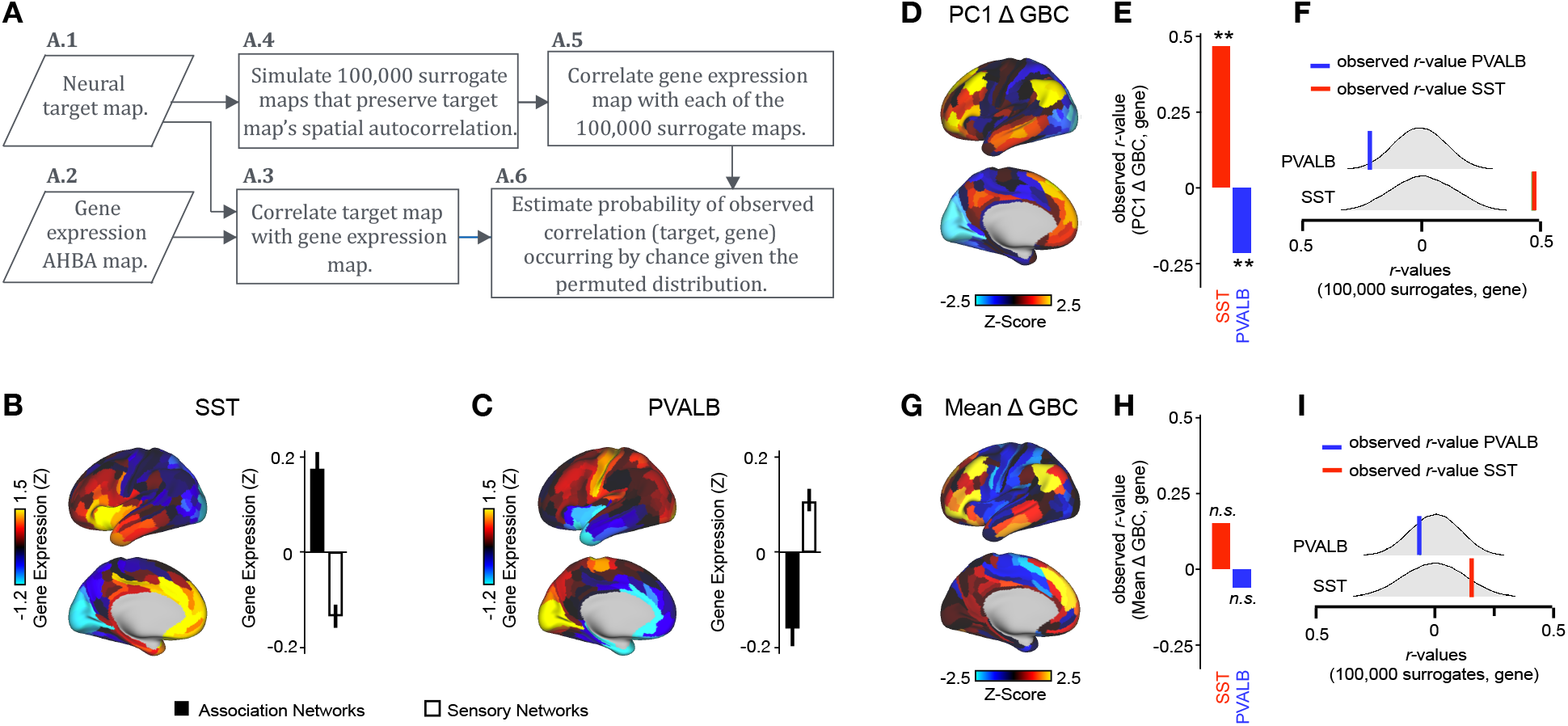
PC1 Δ GBC map tracks SST and PVALB neural gene expression patterns. **(A)** Gene analysis workflow. (A.1) Selection of neural target map. (A.2) Gene expression map was obtained using GEMINI-DOT (50). Specifically, an AHBA gene expression map was obtained using DNA microarrays from six postmortem human brains, capturing gene expression topography across cortical areas. These expression patterns were then mapped onto the cortical surface models derived from the AHBA subjects’ anatomical scans and aligned with the Human Connectome Project (HCP) atlas, described in prior work and Methods (50). (A.3) Correlation of the neural target map with the gene expression map to obtain the observed r value. (A.4) To calculate the significance of the observed r value, we first used BrainSMASH (57) to simulate 100,000 surrogate maps that preserve the neural target map’s spatial autocorrelation. (A.5) We then correlated the gene expression map with each of the 100,000 surrogate maps, to generate a distribution of 100,000 simulated r values. (A.6) Finally, we estimated the probability of the observed correlation between the neural target map and the gene expression map occurring by chance given the permuted distribution. For further details see (**Fig. S7**) **(B)** Gene expression pattern for interneuron marker gene somatostatin (SST). Left: positive (yellow) regions show areas where the gene of interest is highly expressed, whereas negative (blue) regions indicate low expression values. Right: bar plot showing the mean gene expression Z-score across association (black) and sensory (white) networks. **(C)** Gene expression pattern for interneuron marker parvalbumin (PVALB). Left: positive (yellow) regions show areas where the gene of interest is highly expressed, whereas negative (blue) regions indicate low expression values. Right: bar plot showing the mean gene expression Z-score across association (black) and sensory (white) networks. **(D)** Unthresholded PC1 Δ GBC Z-score map (Z-scores computed across 718 parcels). Red/orange areas indicate parcels that have a high positive loading score onto PC1, while blue areas indicate parcels that have a high negative loading score onto PC1. **(E)** Bar plot showing the correlation between PC1 Δ GBC and the following gene expression maps: SST (r=0.47, p<.001) (red) and PVALB (r=−0.22, p=.025) (blue). All p-values are FDR corrected. **(F)** Distribution of 100,000 simulated r values for SST (bottom) and PVALB (top). Bold lines indicate the observed r value between PC1 Δ GBC and SST (red) and PVALB (blue). **(G)** Unthresholded mean Δ (ketamine - placebo) GBC Z-score map at the parcel level (No. parcels = 718) (Z-scores computed across 718 parcels). Red/orange areas indicate regions where participants exhibited stronger GBC in the ketamine condition, whereas blue areas indicate regions where participants exhibited reduced GBC in the ketamine condition, compared with the placebo condition. **(H** Bar plot showing the correlation between mean Δ GBC and the following gene expression maps: SST (r=0.15, p=.363) (red) and PVALB (r=−0.06, p=.627) (blue). All p-values are FDR corrected. **(I)** Distribution of 100,000 simulated r values for SST (bottom) and PVALB (top). Bold lines indicate the observed r value between mean Δ GBC and SST (red) and PVALB (blue).

### Multidimensional behavioral effect of acute ketamine administration

One questions that remains in regards to the PC Δ GBC maps is how the direction of the effect relates to behavior. A principal component is an axis of variation around the mean, and as such direction is uninterpretable. In order to relate ketamine’s behavioral and neural effects, we: (1) data-reduced the behavioral measures; and (2) mapped the data-reduced ketamine-induced changes in behavior to the neural brain effects. We used two types of behavioral measures. The first is an objective measure of cognition collected using a spatial working memory task during the scanner following the resting-state scan. The second is the PANSS, which was used to examine the subjective effects of ketamine retrospectively 180 mins after drug administration. Together these measures resulted in 31 individual items, which are referred to as ‘behavioral measures’ throughout the rest of the paper. We observed significant group mean differences in all behavioral measures: ketamine resulted in impaired cognition and an increase in symptoms in each of the three PANSS subscales (positive, negative, and general symptom scales) (**Fig. 3A**, p<0.05 Bonferroni corrected). For further details see **Table S1**. However, we also observed marked collinearity between behavioral measures, indicating that a dimensionality-reduced solution may capture meaningful variation in participants’ behavioral response to acute ketamine administration (**Fig. 3B**).

**Fig. 3.**
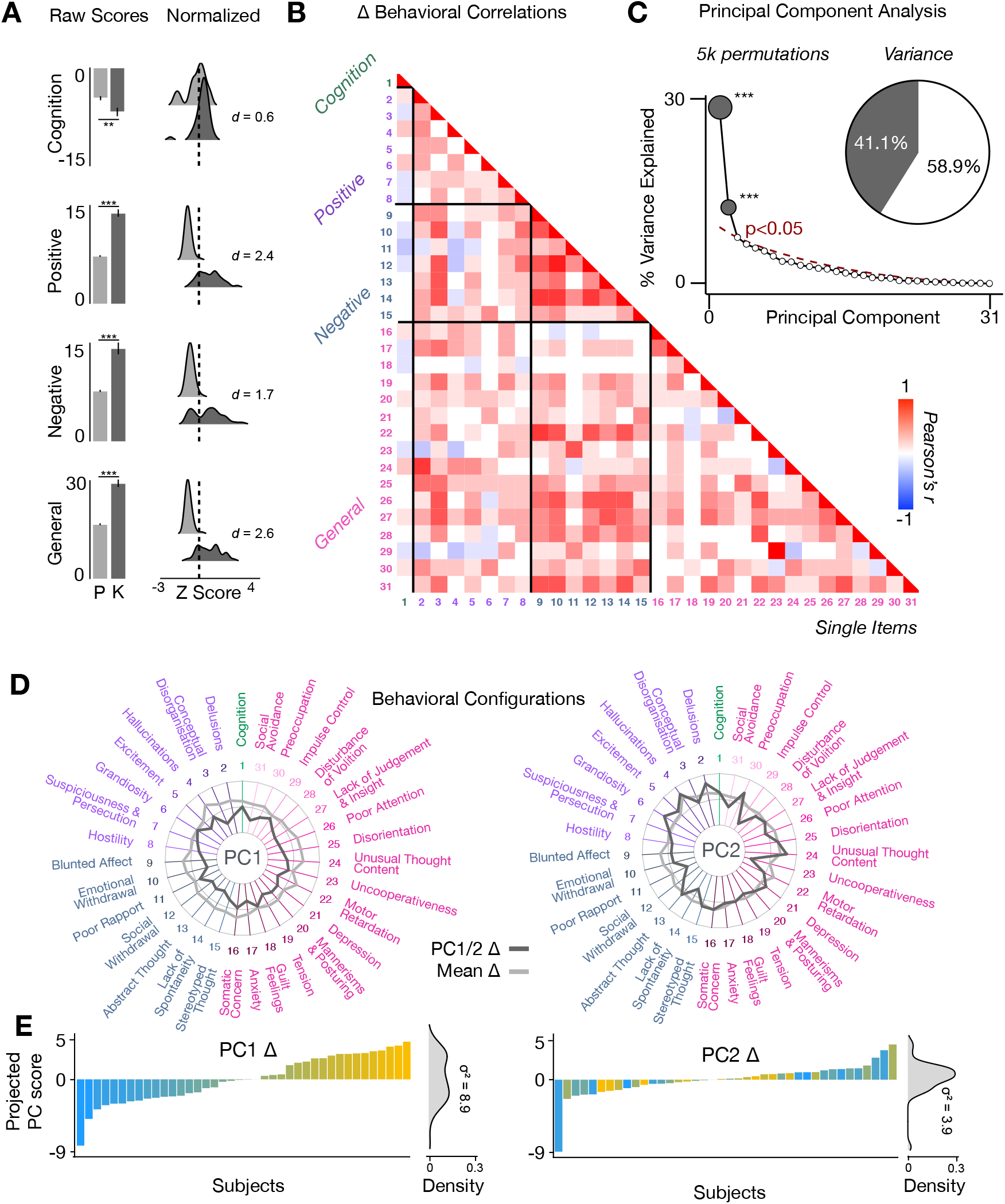
Multidimensional behavioral effect of acute ketamine administration. **(A)** Mean raw scores (left panel) and distribution of normalized scores (right panel) for placebo (P) and ketamine (K) across cognition (spatial working memory) and subjective effects (PANSS positive, PANSS negative, and PANSS general symptoms). Bar plot error bars show standard deviations; distribution plot effect sizes are calculated using Cohen’s d. Condition is colour coded (light grey = placebo, dark grey = ketamine). (** = p<.05, *** = p<.001). **(B)** Correlations between 31 individual item behavioral measures for all participants (N=40). Δ (ketamine - placebo) behavioral measures are used. **(C)** Screeplot showing the % variance explained by each of the principal components (PCs) from a PCA performed using all 31 Δ (ketamine - placebo) behavioral measures across 40 participants. The size of each dark grey point is proportional to the variance explained. The first two PCs (dark grey) survived permutation testing (p<.05, 5000 permutations). Together they capture 41.1% of all symptom variance (inset), with PC1 explaining 29% of the variance and PC2 explaining 12% of the variance. **(D)** Loading profiles shown in dark grey across the 31 behavioral items for PC1 Δ (left) and PC2 (right) Δ. See **Table S3** for numerical values of the behavioral item scores for each PC. The mean Δ score for each behavioral item is also shown in light grey (scaled to fit the same radarplots). Note that the mean Δ configuration resembles the PC1 Δ loading profile more closely than PC2 Δ (which is to be expected as PC1 explains more variance in the behavioral measures). Inner circle = 0.5, middle circle = 0, outer circle = 0.5. **(E)** Bar plot showing the projected PC score for each individual subject (N=40) for PC1 Δ (left) and PC2 (right) Δ. Bars are color coded according to each subject’s PC1 score ranking. The density of the projected PC scores is displayed on the right of each bar plot, demonstrating again that PC1 explains the greatest amount of variance. A paired Pitman-Morgan test revealed that there is a significant difference in variance between the individual subject scores in PC1 Δ and PC2 Δ (t=6.4, df=38, p<.001).

To explore this further, we conducted a data-driven dimensionality reduction procedure of the 31 individual behavioral measures using a PCA. The PCA was computed using the 31 Δ (ketamine - placebo) behavioral measures across all 40 participants. This analysis revealed that ketamine produces two main axes of behavioral alteration, as we identified two significant behavioral PCs that together capture 41.4% of all variance (**Fig. 3C**). This indicates that the behavioral effects of acute ketamine administration are multi-dimensional. The loading configuration of the behavioral measures that form each PC shows that behavioral PC1 indexes variation particularly in the regards to: (1) negative-symptom items such as blunted affect, emotional withdrawal, social withdrawal, abstract thought, and lack of spontaneity; (2) the positive-symptom item conceptual disorganisation; and (3) and general-symptom items such as tension, motor retardation, poor attention, and lack of insight and judgement (**Fig. 3D**). In contrast, behavioral PC2 indexes variation particularly in regards to: (1) the negative-symptom item poor rapport; (2) positive-symptom items such as grandiosity, hallucinations, and delusions; (3) general-symptom items such as uncooperativeness, unusual thought content, impulse control, and preoccupation; and (4) cognition (**Fig. 3D**). See **Table S3** for behavioral PC1 and PC2 symptom values.

### Lower-dimensional behavioral variation reveals robust neuro-behavioral mapping

In order to relate ketamine’s behavioral and neural effects, we then mapped behavioral PC1 and behavioral PC2 to the neural data (52). This resulted in two neuro-behavioral PCs that captured unique patterns of neural variation in comparison to the neural PCA: the neuro-behavioral PC1 map was only moderately correlated with the PC1 Δ GBC map (r=−0.61), while the neuro-behavioral PC2 map was only weakly correlated with PC1-5 Δ GBC (**Fig. S9**). Like PC1 Δ GBC, neuro-behavioral PC1 also differentiates between association and sensory networks (p<.001), and is able to track SST and PVALB gene expression patterns (**Fig. 4A & B, Fig. S10A-C**). We found a significant positive correlation between neuro-behavioral PC1 and SST (r=0.29, p=.009, 100,000 permutations), and trend towards a negative correlation with PVALB (r=−0.14, p=.089, 100,000 permutations) (**Fig. S10A-C**). Meanwhile, though neuro-behavioral PC2 did differentiate between association and sensory networks (p=.04), it did not track SST/PVALB gene expression patterns (**Fig. S10E, Fig. S11E**). Finally, in contrast to PC1-5 Δ GBC maps, the direction of neuro-behavioral PC1-2 is interpretable in relation to behavior. For example, the neuro-behavioral PC1 map demonstrates that a high positive behavioral PC1 score is associated with increased GBC in association networks (e.g. default mode network, fronto-parietal network) and decreased GBC in sensory networks (e.g. secondary visual network), while a high negative behavioral PC1 score is associated with the opposite pattern (**Fig. S11B**).

**Fig. 4.**
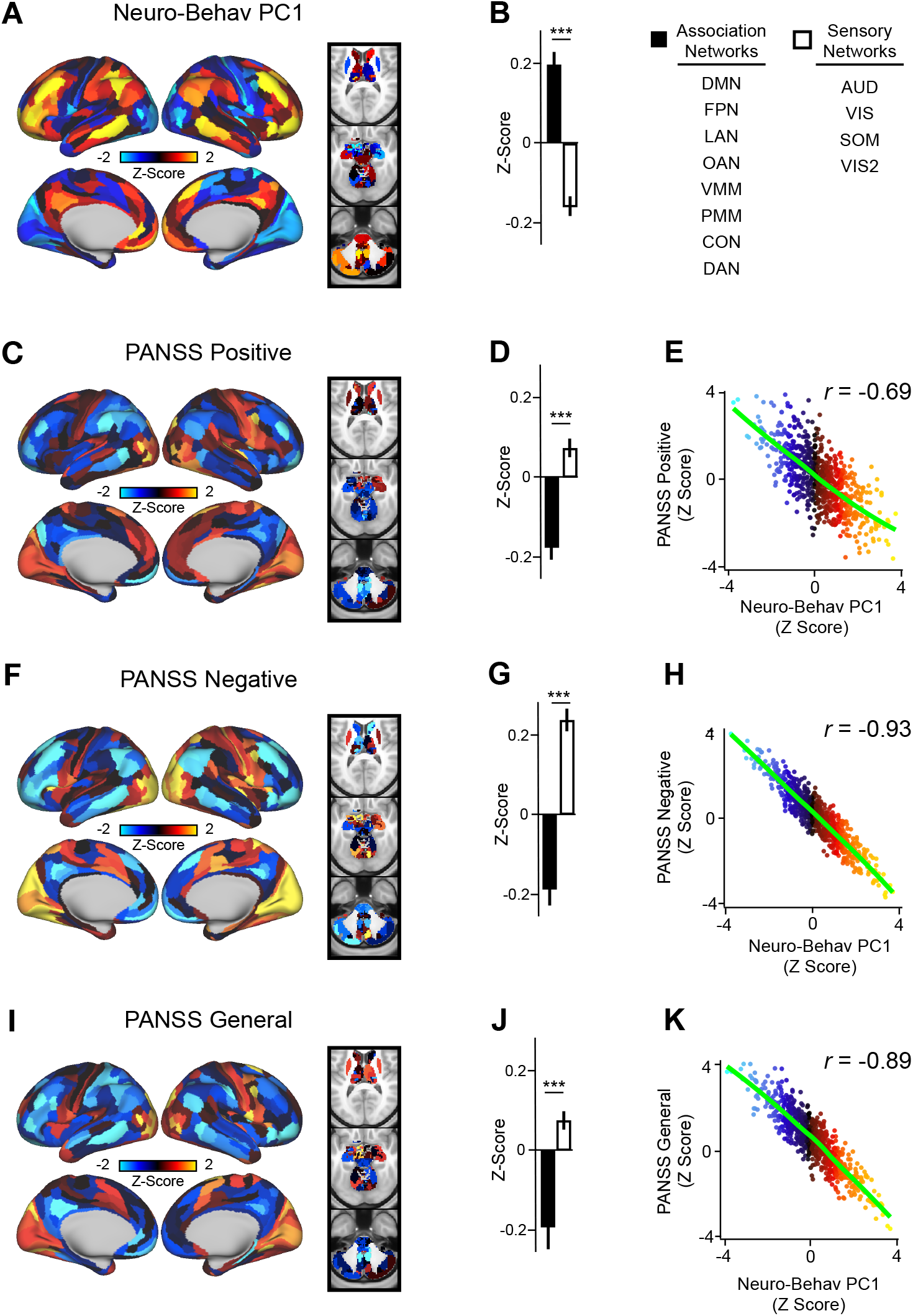
Lower-dimensional behavioral variation reveals robust neuro-behavioral mapping. **(A)** Neuro-behavioral PC1 map showing the relationship between the behavioral PC1 score for each participant regressed onto the Δ GBC map for each participant (N=40). Values shown in each brain parcel are the Z-scored regression coefficient (behavioral PC1 score, Δ GBC) across all 40 subjects. Red/orange areas indicate parcels in which there is a positive relationship between GBC and the behavioral PC1 score, while blue areas indicate parcels in which there is a negative relationship between GBC and the behavioral PC1 score. **(B)** Bar plot showing the mean correlation (Δ GBC, behavioral PC1 score) for association (black) and sensory (white) networks. **(C)** PANSS Positive map showing the relationship between the PANSS Positive score for each participant regressed onto the Δ GBC map for each participant (N=40). Values shown in each brain parcel are the Z-scored regression coefficient (PANSS Positive score, Δ GBC) across all 40 subjects. **(D)** Bar plot showing the mean correlation (Δ GBC, PANSS Positive score) for association (black) and sensory (white) networks. **(E)** Scatterplot showing the relationship across parcels between neuro-behavioral PC1 and PANSS Positive maps (r=0.69, p<.001, n=718). **(F)** PANSS Negative map showing the relationship between the PANSS Negative score for each participant regressed onto the Δ GBC map for each participant (N=40). Values shown in each brain parcel are the Z-scored regression coefficient (PANSS Negative score, Δ GBC) across all 40 subjects. **(G)** Bar plot showing the mean correlation (Δ GBC, PANSS Negative score) for association (black) and sensory (white) networks. **(H)** Scatterplot showing the relationship across parcels between neuro-behavioral PC1 and PANSS Negative maps (r=0.93, p<.001, n=718). **(I)** PANSS General map showing the relationship between the PANSS General score for each participant regressed onto the Δ GBC map for each participant (N=40). Values shown in each brain parcel are the Z-scored regression coefficient (PANSS General score, Δ GBC) across all 40 subjects. **(J)** Bar plot showing the mean correlation (Δ GBC, PANSS General score) for association (black) and sensory (white) networks. **(K)** Scatterplot showing the relationship across parcels between neuro-behavioral PC1 and PANSS General maps (r=0.89, p<.001, n=718).

In order to compare neuro-behavioral PC1 to the established behavioral scales, we also mapped the ketamine-induced changes in PANSS subscales and cognition to the Δ GBC neural brain effects (**Fig. S12**). We found that the PANSS Negative and General neural maps were both highly negatively correlated with the the neuro-behavioral PC1 map, while the PANSS Positive map was moderately correlated with the neuro-behavioral PC1 map, and the cognition map showed no correlation (**Fig. 4**). In contrast, neither the PANSS subscales nor cognition showed a strong correlation with neuro-behavioral PC2, though there was a moderate positive correlation between PANSS Positive and neuro-behavioral PC2 (r=0.42) (**Fig. S13B**). In addition, we found that a similar amount of variation was captured by neuro-behavioral PC1 (*σ*^2^ = 0.07), PANSS Negative (*σ*^2^ = 0.08), and PANSS General (*σ*^2^ = 0.05) (**Fig. S14B, H & J**). We also showed that as with neuro-behavioral PC1, PANSS Negative and PANSS General were all able to successfully track SST and PVALB neural gene expression patterns (**Fig. S10**). PANSS Negative was significantly negatively correlated with SST (r=−0.34, p<.001, 100,000 permutations) and significantly positively correlated with PVALB (r=0.16, p=.05, 100,000 permutations) (**Fig. S10J-L**). PANSS General was also significantly negatively correlated with SST (r=−0.26, p=.019, 100,000 permutations) and significantly positively correlated with PVALB (r=0.18, p=.04, 100,000 permutations) (**Fig. S10M-O**). Finally, as the PANSS literature has indicated that a five-factor model may be more stable than the original three-factor model, we also repeated this analysis using the following five PANSS factors: Positive, Negative, Disorganization, Excitement, and Emotional Distress (see **Fig. S15**) (58). Overall, the 3-factor and 5-factor PANSS yielded similar results, though neuro-behavioral PC2 showed a much stronger positive correlation with the 5-factor PANSS Positive map (r=0.73) than it did with the 3-factor PANSS Positive map (r=0.42) (**Fig. S13B, Fig. S15C**).

### Comparing individual variation in ketamine-induced behavioral and neuro-behavioral changes

In order to compare individual variation in ketamine-induced behavioral and neuro-behavioral changes, we first established that there is a consistent effect, at the individual subject level, between the behavioral and neural effects of ketamine along the data-driven dimensions. We hypothesized that subjects with high behavioral scores along a particular dimension, such as PC1, should also exhibit strong neural similarity to the neuro-behavioral PC1 map. To test this, we first rank-ordered each individual subject’s behavioral PC1 score (**Fig. 5A**). The highest negative loader (subject 22) had a behavioral PC1 score of −2.8, while the highest positive loader (subject 19) had a score of 1.6. We then plotted each individual subject’s neuro-behavioral PC1 score (**Fig. 5B**). Here, the neuro-behavioral PC1 score was calculated by correlating each individual subject’s Δ GBC map with the neuro-behavioral PC1 map (**Fig. 4A**). This provides an index of the similarity between the individual subject’s neural GBC and the map of group-level neural changes relevant to the PC1 behavioral dimension. As with the behavioral effect, subject 22 was the highest negative loader (score = −2.4), whereas subject 19 ranked as the fifth highest neuro-behavioral positive loader (score = 1.2) (**Fig. 5A & B**). Furthermore, we found that the top five highest neuro-behavioral positive loaders’ individual Δ GBC map were all significantly correlated with each other (r values ranging from 0.16 to 0.35) (**S16**). Finally, we correlated behavioral PC1 scores with neuro-behavioral PC1 scores, which demonstrated that the two measures are highly positively correlated (r=0.7, p<.001) (**Fig. 5C**). Overall, this demonstrates that the strength of behavioral and neural ketamine-induced changes along the PC1 dimension appear to be largely consistent across individuals, i.e. individuals with a strong behavioral effect will also show the greatest change in GBC within behaviorally-relevant neural circuits.

**Fig. 5.**
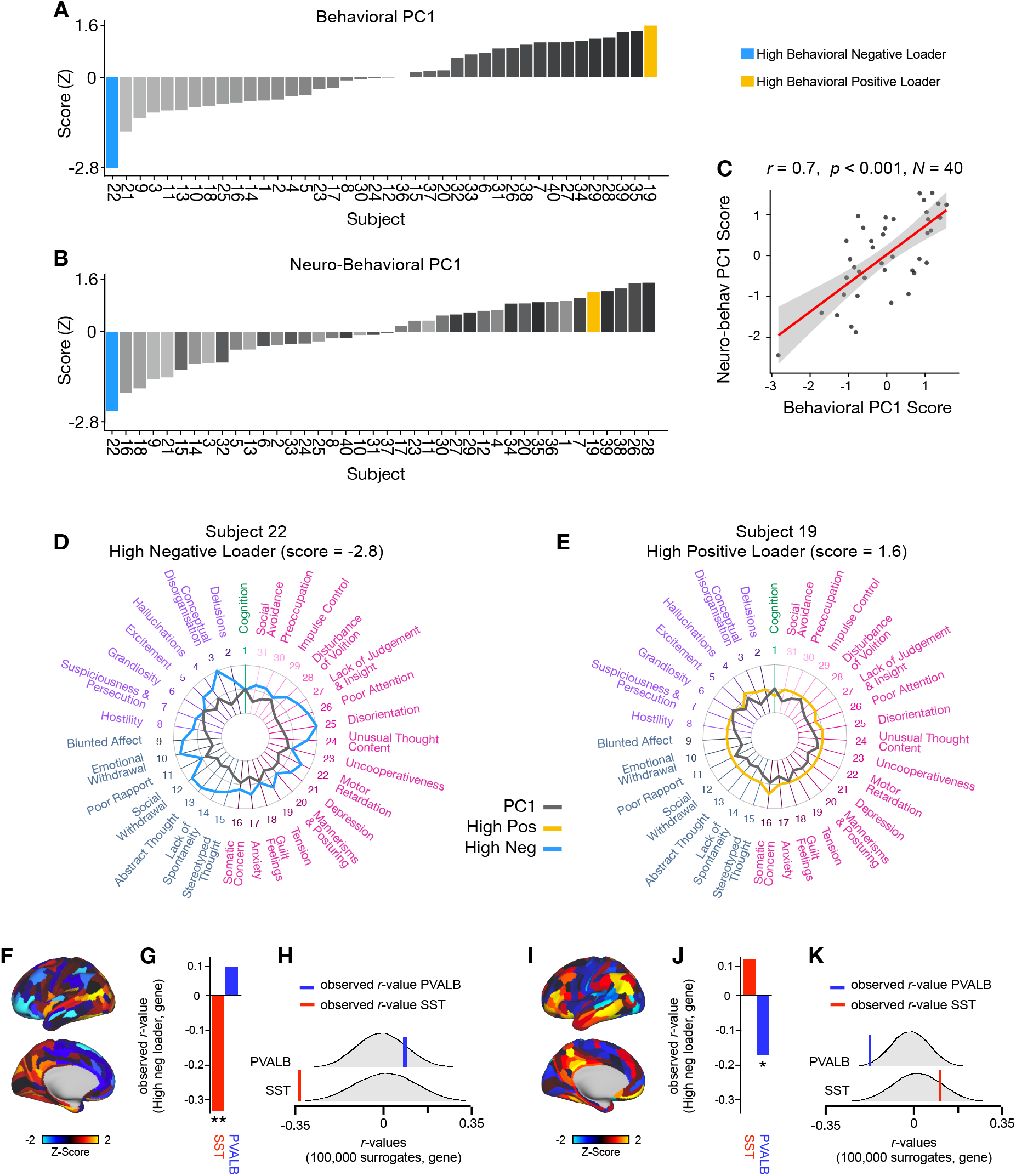
Individual variation in ketamine-induced behavioral and neuro-behavioral changes. **(A)** Bar plot showing the behavioral PC1 score (Z) for each individual participant (N=40). Bars are ordered, numbered, and color-coded according to each participant’s behavioral PC1 score (light grey = highly negative score, dark gray = highly positive score). Blue = high negative behavioral PC1 loader, yellow = high positive behavioral PC1 loader. **(B)** Bar plot showing the neuro-behavioral PC1 score (Z) for each individual participant (N=40). The neuro-behavioral PC1 score is calculated by correlating each participant’s Δ GBC map with the neuro-behavioral PC1 map, and then Z-scoring the r-values. Bars are ordered according to each participant’s neuro-behavioral PC1 score, but labelled and color-coded according to each participant’s behavioral PC1 score (light grey = highly negative score, dark gray = highly positive score). Blue = high negative behavioral PC1 loader, yellow = high positive behavioral PC1 loader. **(C)** Scatterplot showing the correlation between each participant’s behavioral PC1 score and their neuro-behavioral PC1 score. Overall participants behavioral PC1 score and neuro-behavioral PC1 score were highly correlated (r=0.7, p<.001). **(D & E)** Loading profile (raw scores) shown in blue across the 31 behavioral items for subject 22, who had the highest negative behavioral PC1 score (left); and subject 19, who had the highest positive behavioral score (right). The behavioral PC1 score for each behavioral item is also shown in dark grey (scaled to fit the same radarplots). Inner circle = 0.5, middle circle = 0, outer circle = 0.5. **(F)** Δ GBC map for subject 22. **(G)** Bar plot showing the correlation between subject 22 Δ GBC and the following gene expression maps: SST (r=−0.33, p<.001) (red) and PVALB (r=0.08, p=.21) (blue). All p-values are FDR corrected. **(H)** Distribution of 100,000 simulated r values for subject 22 Δ GBC and SST (bottom) and subject 22 Δ GBC and PVALB (top). Bold lines indicate the observed r value between PC1 Δ GBC and SST (red) and PVALB (blue). **(I)** Δ GBC map for subject 19. **(J)** Bar plot showing the correlation between subject 19 Δ GBC and the following gene expression maps: SST (r=0.1, p=.21) (red) and PVALB (r=−0.17, p=.027) (blue). All p-values are FDR corrected. **(K)** Distribution of 100,000 simulated r values for subject 19 Δ GBC and SST (bottom), and subject 19 Δ GBC and PVALB (top). Bold lines indicate the observed r value between PC1 Δ GBC and SST (red) and PVALB (blue).

Next, to connect across behavior, neural, and molecular mechanisms at the single subject level, we explored whether individual subjects’ Δ GBC maps could also track SST and PVALB neural gene expression topography. Here, we hypothesized that the highest positive and negative behavioral PC1 loaders’ Δ GBC maps would correlate in opposing directions with the maps of SST and PVALB expression, thus providing insight into which distinct circuit-level mechanisms may underlie the individual differences in systems-level neural and behavioral effects of ketamine. We first focused on the highest negative behavioral PC1 loader, subject 22. Following acute ketamine administration, subject 22 experienced increased conceptual disorganisation, emotional withdrawal, social withdrawal, abstract thought, poor attention, and disorientation (**Fig. 5D**). We then examined subject 22’s Δ GBC and found that they showed hypoconnectivity across association network parcels and hyperconnectivity across sensory network parcels (**Fig. 5F**). In addition, there was a significant difference between connectivity in association and sensory networks (t=−4.99, df=1151, p<.001). Lastly, we explored whether subject 22’s Δ GBC would track SST and PVALB neural gene expression topography. We found a significant negative correlation between subject 22’s Δ GBC and the SST gene expression map (r=−0.33, p<.001, 100,000 permutations) well beyond what is expected by chance alone (**Fig. 5G-H**). However, no relationship was found with the PVALB gene expression map (r=0.08, p=.21, 100,000 permutations).

We then repeated the same analysis for the highest positive behavioral PC1 loader, subject 19. Following acute ketamine administration, subject 19 experienced a reduction in cognition and excitement, improved attention, and no change in most other symptoms (**Fig. 5E**). Subject 19’s Δ GBC map showed hyperconnectivity across association network parcels and hypoconnectivity across sensory network parcels (**Fig. 5I**). In addition, there was a significant difference between connectivity in association and sensory network parcels (t=5.68, df=1367, p<.001). Lastly, we explored whether subject 19’s Δ GBC would track SST and PVALB neural gene expression topography. We found a significant negative correlation between subject 19’s Δ GBC and the PVALB gene expression map (r=−0.17, p=.027, 100,000 permutations) well beyond what is expected by chance alone (**Fig. 5J-K**). However, no relationship was found with the SST gene expression map (r=0.1, p=.21, 100,000 permutations). Collectively, these results provide insight into the differentiating mechanisms, modulated by somatostatin and parvalbumin interneurons, which may underlie individual variation in acute responses to ketamine. We demonstrate that the data-driven axis of behavioral variation provides an anchor point for neural Δ GBC effects and molecular mechanisms that are interpretable at the single subject level.

## Discussion

Ketamine has emerged as one of the most promising treatments for treatment-resistant depression (1, 4). However, two key knowledge gaps remain. First of all, inter-individual variability in response to ketamine is still not well understood. Secondly, it is unclear how ketamine’s downstream molecular mechanisms connect to its neural and behavioral effects. The current study addresses these knowledge gaps by showing that: (1) both the neural and behavioral effects of acute ketamine are multi-dimensional, reflecting substantial inter-individual variability in response to ketamine; (2) the use of a multi-dimensional neural approach provides insight into the downstream molecular mechanisms of ketamine’s neural effects, specifically in regards to SST and PVALB interneurons; and (3) the use of a multi-dimensional behavioral approach provides an anchor point for both ketamine’s neural effects and molecular mechanisms at the single-subject level.

### Ketamine’s neural effects are multi-dimensional, reflecting inter-individual variability

We show that the neural geometry of ketamine is multi-dimensional, as ketamine results in five bi-directional neural axes of variation (**Fig. 1A, Fig. S3**). Interpreting this finding is of particular interest given that ketamine is a molecule that acts at the same target in every human brain, antagonising the NMDA receptor, which should result in a single uni-directional principal component (PC). However, we found ketamine results in multiple bi-directional neural PCs (**Fig. 1, Fig. S3**). The bi-directional nature of the five neural PCs highlights the inter-individual variability in response to ketamine, as extreme positive and negative PC1 Δ GBC loaders show the opposite pattern of change in GBC in numerous brain regions following ketamine. For example, high positive PC1 Δ GBC loaders exhibit increased GBC in association networks such as the default mode network, and decreased GBC in sensory cortices such as the secondary visual network following ketamine, while high negative PC1 Δ GBC loaders show the reverse pattern (**Fig. S8**). The bi-directional nature of the PCs helps explain the contradictory results in the ketamine resting-state literature as the the direction of the effects are often inconsistent (10–12, 15, 19).

A number of possible interpretations exist for why ketamine results in multiple bi-directional neural PCs. For example, one PC may reflect ketamine’s effects in excitatory-excitatory (E-E) synapses, while the another may reflect ketamine’s effects in excitatory-inhibitory (E-I) synapses. Another possibility is that given we know ketamine’s affinity for certain cells is based on their NMDA receptor subunit configuration, one PC may reflect ketamine’s ‘high affinity’ actions on GluN2C and GluN2D containing receptors, while another PC may reflect ketamine’s ‘lower affinity’ actions on GluN2A and GluN2B containing receptors (41**?**, 42). A third possibility is that the multiple PCs may be driven by ketamine’s numerous microcircuit level targets, as ketamine is known to effect: (1) SST interneurons, which in turn disinhibit glutamatergic pyramidal distal dendrites and PV neurons; (2) PV neurons, which disinhibit the soma of glutamatergic pyramidal neurons; (3) VIP neurons, which may reduce distal dendrite inhibition but disinhibit SST activity; and (4) chandelier cells, which permit the back-propagation of action potentials from the axon to the soma ().

Ketamine’s numerous microcircuit level targets may also explain why ketamine shows increased dimensionality compared to psilocybin and LSD (**Fig. 1, Fig. S6**). In comparison to ketamine, psilocybin and LSD have fewer microcircuit level targets: both psilocybin and LSD’s behavioral effects are predominantly linked to 5HT2A agonism, and 5HT2A receptors are located on fewer cellular elements (59–62). In addition, both psilocybin and LSD are excitatory while ketamine results in increased excitation indirectly as a consequence of the inhibition of interneurons (37, 63). Ketamine’s multi-dimensional neural effects may be due to complementary inhibitory and excitatory dimensions which are not present to the same extent with psilocybin and LSD.

### Ketamine’s data-driven principal neural gradient captures its hypothesized molecular mechanisms better than the mean effect

We found ketamine’s principal neural gradient of change in GBC captures its hypothesized molecular mechanisms better than the mean effect, as did PC3-5 Δ GBC (**Fig. 2, Fig. S5**). For example, ketamine’s principal neural gradient exhibited a strong positive relationship with SST gene expression and a strong negative relationship with PVALB gene expression well beyond chance alone (**Fig. 2E**). In contrast, no significant relationship was found between ketamine’s mean effect and SST/PVALB gene expression maps (**Fig. 2H**). This finding extends previous studies by demonstrating that ketamine’s primary axis of neural variation may be attributable to the stimulation of SST and PVALB interneurons, providing further support for the indirect ketamine hypothesis (37). Furthermore, as PC1 Δ GBC is a bidirectional axis of variation, it indicates that in some individuals ketamine-induced neural effects will exhibit a strong relationship with SST, while in others ketamine-induced neural effects will show a strong relationship with PVALB (**Fig. S8**). Again, there are a number of possible explanations for this inter-individual variability. For instance, previous literature has demonstrated that SST and PVALB gene expression varies across individuals and is altered by stress, drugs, and psychiatric diagnoses (64, 65). Overall, given that the relationship between ketamine’s hypothesized SST and PVALB molecular mechanisms of action and its neural effects is only evident when using a multi-dimensional approach, it is important future ketamine studies consider a multi-dimensional rather than mean-based approach.

### Ketamine results in two principal behavioral gradients which capture novel neural variation

As with ketamine’s neural geometry, we found that ketamine’s behavioral geometry was also multi-dimensional: ketamine resulted in two bi-directional behavioral axes of variation (**Fig. 3**). Again, this demonstrates that there is inter-individual variation in the behavioral response to ketamine that is not fully captured by the mean. For example, while behavioral PC1 is closely related to the mean effect, behavioral PC2 tracks changes in ‘uncooperativeness’, ‘impulse control’, and ‘poor rapport’ that are not captured by the mean effect (**Fig. 3D**). This finding is also interesting in relation to the wider literature. Although PANSS items are known to show substantial collinearity between symptom measures, most data-reduction studies have identified a five-factor solution rather than the two-factor solution we observe here (58). While it may be that our sample size lacks adequate power to detect additional factors, previous PANSS data-reduction studies have focused on psychosis populations rather than acute ketamine administration.

These two behavioral PCs are also of interest as when they are regressed onto the neural Δ GBC data, the two resulting neuro-behavioral PC maps: (1) capture novel patterns of neural variation; (2) have a directionality which is interpretable in relation to behavior (unlike the PC1-5 Δ GBC); (3) show a close relationship to both the 3-factor and 5-factor PANSS subscales; and (4) track SST and PVALB gene expression patterns (**Fig. 4, Fig. S10**). The novel neural variation captured by the neuro-behavioral PCs is evidenced by the fact that although neuro-behavioral PC1 is moderately correlated with PC1 Δ GBC (r=−0.61), neuro-behavioral PC2 and PC2 Δ GBC were only weakly correlated (r=−0.12) (**Fig. S9**). In contrast to PC1-5 Δ GBC, the neuro-behavioral PCs are interpretable in relation to behavior. Such an understanding is crucial for informing predictions about how a specific individual will respond to ketamine in a clinical setting. Importantly, this prediction may ultimately be done using only their symptom profile, without the need for costly neural scans.

When we directly compared our data-driven approach to the existing 3-factor and 5-factor PANSS scales, we found the 3-factor and 5-factor approach were both sufficient to explain neuro-behavioral PC1. For example, as 3-factor PANSS Negative, 3-factor PANSS General, 5-factor PANSS Negative, and 5-factor PANSS Disorganization were all highly correlated with neuro-behavioral PC1 (**Fig. S13C & E, Fig. S15E & H**). In contrast the 5-factor model was more closely related to neuro-behavioral PC2, as 5-factor PANSS Positive was highly correlated with neuro-behavioral PC2 while none of the 3-factor PANSS subscales were highly correlated with neuro-behavioral PC2 (**Fig. S13, Fig. S15C**). Finally, we found that neuro-behavioral PC1, PANSS Negative, and PANSS General all tracked SST and PVALB gene expression patterns, allowing us to relate behavior to both neural and molecular mechanisms (**Fig. S10B, J & K**). This speaks to the validity of the existing PANSS subscales, though our results indicate that a 2-factor rather than a 3-factor or 5-factor solution may be optimal for detecting the effects of acute ketamine administration.

### Ketamine’s data-driven principal behavioral gradient provides an anchor point for neural effects and molecular mechanisms at the single subject level

In terms of developing a framework for connecting across ketamine’s behavioral, neural, and molecular effects, the ultimate goal is to be able take an individual’s behavioral response to a molecular perturbation, and precisely predict their imaging effect, or vice versa. When we tested our neuro-behavioral model by comparing the order in which individual participants load onto behavioral PC1 and neuro-behavioral PC1, we found the order was largely preserved (**Fig. 5A-C**). Thus, this framework enables us to take an individual’s behavioral response to ketamine, and then meaningfully infer where that individual falls in relation to their ketamine induced neural effects. The fact that this framework operates at the whole-brain level indicates that the neural effect of ketamine is not an ROI-based effect but a distributed system-wide effect. Furthermore, we found that both the highest positive and negative behavioral PC1 loaders’ Δ GBC maps track SST and PVALB gene expression patterns (**Fig. 5F-K**). By providing a framework with which to connect across ketamine’s behavioral, neural and molecular effects at the individual subject level, this study may also have implications for the development of new treatments. The identification of the targets responsible for the different behavioral effects of ketamine, and potentially its metabolites, is critical for the development of novel pharmaco-therapies that lack the side effects of ketamine such as the psychotomimetic effects, changes in sensory perception, and abuse potential.

### Limitations and future directions

As with most psychoactive pharmacological neuroimaging studies, participants were able to correctly identify whether they received the substance or saline 100% of the time, even with a double-blind design. Future studies may be able to mitigate this by using an active placebo control. In addition, as the ketamine and saline scans took place on the same day, ketamine’s residual effects ruled out the counterbalancing of the order; furthermore, the study design did not collect follow-up data on post-scan days, which may be of interest given that the antidepressive effects of ketamine are shown to peak 24-72 hours post-infusion (1, 66). Future studies may wish to address these limitations by spacing out the scans and collecting additional scans on the day after ketamine infusion. The present study used GBC: a dimension-reduced summary measure of neural resting-state fMRI. While this remains a principled method of reducing the feature space, it is possible that some of the behaviorally relevant neural information may be lost by first summarizing neural functional connectivity features in this manner. Thus, future studies may wish to explore the multi-dimensional effects of ketamine using the full neural functional connectivity matrix for each subject. In this analysis, we largely focused on PC1-2 Δ GBC when exploring ketamine’s neural effects as the bulk of the neural variance was explained by PC1 and PC2 Δ GBC. To unpack the variation in PC3-5 Δ GBC a larger sample size is needed, especially given each PC represents a bidirectional axis of variation. A larger sample size would also allow us to investigate alternative models for ketamine’s molecular mechanisms in addition to SST and PVALB, such as how ketamine’s effects differ based on NMAR subtypes, or excitatory vs. inhibitory synapses. Finally, future studies may also wish to examine intra-individual variation in addition to inter-individual variation in response to ketamine.

## Conclusions

A key goal in psychiatric research is to predict treatment response. In relation to ketamine, two key barriers to the development of predictive biomarkers for treatment response remain. First of all, inter-individual variability in response to ketamine is still not well understood. Secondly, it is unclear how ketamine’s downstream molecular mechanisms connect to its neural and behavioral effects. We address both these knowledge gaps by: (1) showing that there is substantial inter-individual variability in both the behavioral and neural response to ketamine that requires a multi-dimensional approach and is not sufficiently captured by the mean; and (2) providing a multi-dimensional framework with which to connect across ketamine’s behavioral, neural and molecular effects at the individual subject level. This multi-dimensional framework has the potential to generate predictions about how an individual will respond to ketamine at the individual subject level. It is of paramount importance for future studies to test this in patients using actual treatment response data.

## ACKNOWLEDGEMENTS

We would like to thank Dr. Robert Malison for his contribution to the conceptualization of the study and analyses. We would also like to thank the Yale Magnetic Resonance Research Center (MRRC) and the Yale Center for Clinical Investigation (YCCI). This study was supported by NIH grants DP5OD012109-01 (A.A.), 1U01MH121766 (A.A.), R01MH112746 (J.D.M.), 5R01MH112189 (A.A.), 5R01MH108590 (A.A.), NIAAA grant 2P50AA012870-11 (A.A.); NSF NeuroNex grant 2015276 (J.D.M.); Brain and Behavior Research Foundation Young Investigator Award (A.A.); SFARI Pilot Award (J.D.M., A.A.); Heffter Research Institute (Grant No. 1–190420); Swiss Neuromatrix Foundation (Grant No. 2016–0111m Grant No. 2015 – 010); Swiss National Science Foundation under the framework of Neuron Cofund (Grant No. 01EW1908), Usona Institute (2015 – 2056), ARRS grants P3-0338 (G.R.), J7-8275 (G.R.).

## Methods

### Ketamine Study Participants

Healthy participants were recruited from the New Haven area via flyers and online ads. In order to be eligible to receive ketamine, participants were required to meet the following set of criteria, as determined by a detailed telephone interview and an in-person clinical assessment: i) Age 21-60; ii) IQ>70 as measured via Wide Range Achievement Test (WRAT-3) and the Wechsler Adult Intelligence Scale (WAIS-III); iii) intact or corrected-to-normal vision; iv) weight < 300 lbs.; v) MR safe (free of metallic objects and absence of claustrophobia); vi) no serious medical or physical conditions, as confirmed by a self-report, electrocardiogram, blood work, and physical examination by a licensed physician; vii) no lifetime neurological or psychiatric diagnoses; viii) minimal alcohol intake and no use of psychoactive drugs or history of abuse/dependence, confirmed by interview and urinalysis; ix) no first-degree relatives with DSM Axis-I diagnoses or alcohol/substance abuse history; x) no known sensitivity to ketamine or heparin; xi) no donation of blood in excess of 500ml within 2 months of participation. Eligible subjects provided informed consent approved by Yale University Institutional Review Board. Forty participants took part in the study. The sample size was determined ahead of study initiation based on prior studies (24, 67–70), which would achieve statistical power of >86% for a estimated medium effect size (Cohen’s d of 0.5). Demographic details can be found in **Table S2**).

### Ketamine Study Design

The protocol was approved by the Yale Human Investigations Committee (ClinicalTrials.gov Identifier: NCT03842800). All subjects provided written informed consent. The study employed a double-blind within-subjects design. Placebo was administered during the first neuroimaging scan session, and ketamine (initial bolus 0.23 mg/kg, continuous infusion 0.58 mg/kg/hour) during the second, as residual ketamine effects ruled out counterbalancing of the order. Participants were not informed about the order in which they would receive saline and ketamine, however they were able to correctly identify the ketamine infusion 100% of the time. Cognitive effects were measured using a Spatial Working Memory task completed in the scanner. Subjective effects were measured before and after the scan (180 minutes post drug administration) using the following scales: (1) Positive and Negative Syndrome Scale (PANSS), (2) Clinician Administered Dissociative States Scale (CADSS), and (3) Beck’s Depression Inventory (BDI).

### Ketamine Infusion Protocol

Participants were instructed to fast for 12 hours prior to the scan and avoid any alcohol or medications 72 hours prior to the scan. As a precaution, blood alcohol content was assessed with an electronic breathalyzer on the morning of the infusion. Prior to entering the scanner, subjects received IV cannulation in each forearm: one IV for the placebo/ketamine infusion and one for blood draws during the scan. In the first scan session, an initial bolus of saline was administered over two minutes prior to collecting BOLD images, followed by a continuous maintenance infusion of saline throughout the remainder of the scan. In the second scan session, an initial bolus of racemic ketamine (0.23 mg/kg) diluted in saline solution was delivered just prior to data acquisition, followed by a continuous infusion of racemic ketamine (0.58 mg/kg/hour) throughout the remainder of the scan. Blood was drawn immediately after the resting-state run, and subsequent gas chromatography-mass spectrometry determined that the group mean serum ketamine concentration at the end of this run was 470 nmol/L. The scan was aborted and the participant removed for evaluation if, at any point, they expressed discomfort related to administration of the drug or placebo, or if they became unresponsive or showed any indications of an adverse reaction. Diazepam was available for anxiety in the event of a particularly negative response, but was never needed. Full neuroimaging protocol details can be found in **Fig. S17**.

### Ketamine Neuroimaging Data Acquisition

Neural data were collected using a Siemens 3T scanner with with a 32 channel head coil at the Yale Center for Biomedical Imaging. Imaging acquisition parameters were aligned with those of the Human Connectome Project (HCP) (71). High-resolution T1w and T2w structural images were acquired in 224 AC-PC aligned slices, 0.8mm isotropic voxels. T1w images were collected with a magnetization-prepared rapid gradient-echo (MP-RAGE) pulse sequence (time repetition (TR) = 2400 ms, time echo (TE) = 2.07 ms, flip angle = 8°, field of view = 256 × 256 mm). T2w images were collected with a SCP pulse sequence (Tr=3200 ms, TE = 564 ms, flip angle = T2 var, field of view = 256 × 256 mm). Resting-state BOLD images were collected with a multi-band accelerated fast gradient-echo, echo-planar sequence (acceleration factor=6, Tr=700 ms, TE = 31.0 ms, flip angle = 55°, field of view = 210 × 210 mm, matrix = 84 × 84, bandwidth = 2290 Hz); 54 interleaved axial slices aligned to the anterior-posterior commissure (AC-PC) with 2.5mm isotropic voxels. 400 volumes were acquired per resting state scan resulting in a scan duration of 4.67 minutes. Additionally, a pair of reverse phase-encoded spin-echo field maps (anterior-to-posterior and posterior-to-anterior) were acquired (voxel size = 2.5 mm isotropic, Tr=7220 ms, TE = 73 ms, flip angle = 90°, field of view = 210 × 210 mm, bandwidth = 2290 Hz).

### LSD and Psilocybin Neuroimaging Data Acquisition

To assess whether increased dimensionality is specific to ketamine we compared the neural effects of ketamine to the neural effects of lysergic acid diethylamide (LSD) and psilocybin. Methods for the LSD neuroimaging study (N=24) are described in detail in prior publications (51). The use of LSD in humans was authorized by the Swiss Federal Office of Public Health, Bern, Switzerland. The study protocol was approved by the Cantonal Ethics Committee of Zurich (KEK-ZH_No: 2014_0496). The study employed a fully double-blind, randomized, within-subject cross-over design with three conditions: (1) placebo + placebo condition: placebo (179 mg Mannitol and Aerosil 1 mg po) after pretreatment with placebo (179 mg Mannitol and Aerosil 1 mg po); (2) placebo + LSD condition: LSD (100 μg po) after pretreatment with placebo (179 mg Mannitol and Aerosil 1 mg po), or (3) Ketanserin + LSD condition: LSD (100 μg po) after pretreatment with the 5-HT2A antagonist Ketanserin (40 mg po). Data were collected for all subjects in a randomized counterbalanced order at three different sessions each two weeks apart. For all conditions, the first substance was administered 60 min before the second substance, and the first neural scan was conducted 75 min after the second administration, with a second scan conducted at 300 min post-administration. In the present study, only data from the (1) placebo + placebo and (2) placebo + LSD conditions were evaluated. Methods for the psilocybin neuroimaging study (N=23) are described in detail in prior publications (53). The use of psilocybin in humans was authorized by the Swiss Federal Office of Public Health, Bern, Switzerland. The study was registered at ClinicalTrials.gov (NCT03736980). The study employed a fully double-blind, randomized, within-subject cross-over design. Participants at 2 different occasions 2 weeks apart received either placebo (179 mg mannitol and colloidal silicon dioxide [Aerosil; Evonik Resource Efficiency GmbH, Essen, Germany] 1 mg orally; placebo condition) or psilocybin (0.2 mg/kg orally; psilocybin condition). The resting-state scan was conducted at 3 time points between administration and peak effects: 20, 40, and 70 minutes after treatment administration. In the present study, only data from the psilocybin and placebo neural scans conducted at 70 minutes were evaluated.

### Neuroimaging Data Preprocessing

Neuroimaging data was preprocessed using the Human Connectome Project (HCP) minimal preprocessing pipeline (72). A summary of the HCP pipelines is as follows. First T1/2-weighted structural images were corrected for bias-field distortions and then warped to the standard Montreal Neurological Institute-152 (MNI-152) brain template in a single step, through a combination of linear and non-linear transformations via the FM-RIB Software Library (FSL) linear image registration tool (FLIRT) and non-linear image registration tool (FNIRT) (73). Next, FreeSurfer’s recon-all pipeline was used to segment brain-wide gray and white matter to produce individual cortical and subcortical anatomical segmentations (74). Cortical surface models were generated for pial and white matter boundaries as well as segmentation masks for each subcortical grey matter voxel. Using the pial and white matter surface boundaries, a ‘cortical ribbon’ was defined along with corresponding subcortical voxels, which were combined to generate the neural file in the Connectivity Informatics Technology Initiative (CIFTI) volume/surface ‘grayordinate’ space for each individual subject (72). BOLD data were motion-corrected by aligning to the middle frame of every run via FLIRT in the initial NIFTI volume space. In turn, a brain-mask was applied to exclude signal from non-brain tissue. Next, cortical BOLD data were converted to the CIFTI gray matter matrix by sampling from the anatomically-defined gray matter cortical ribbon and subsequently aligned to the HCP atlas using surface-based nonlinear deformation (72). Subcortical voxels were aligned to the MNI-152 atlas using whole-brain non-linear registration and then the Freesurfer-defined subcortical segmentation applied to isolate the sub-cortical grayordinate portion of the CIFTI space.

After the HCP minimal preprocessing pipelines, movement scrubbing was performed (75). As in prior work (3), all BOLD image frames with possible movement-induced artifactual fluctuations in intensity were flagged using two criteria: frame displacement (the sum of the displacement across all six rigid body movement correction parameters) exceeding 0.5 mm (assuming 50 mm cortical sphere radius) and/or the normalized root mean square (RMS) (calculated as the RMS of differences in intensity between the current and preceding frame, computed across all voxels and divided by the mean intensity) exceeding 1.6 times the median across scans. Any frame that met one or both of these criteria, as well as the frame immediately preceding and immediately following, were discarded from further preprocessing and analyses. Subjects with more than 50% frames flagged using these criteria were excluded. Next, a high-pass filter (threshold 0.008 Hz) was applied to the BOLD data to remove low frequency signals due to scanner drift. In-house Matlab code was used to calculate the average variation in BOLD signal in the ventricles, deep white matter, and across the whole grey matter (’global signal’), as well as movement parameters. These signals, as well as their first derivatives to account for delayed effects, were then regressed out of the grey matter BOLD time series as nuisance variables (as any change in the BOLD signal due to these variables are not likely to reflect neural activity) (76). It should be noted that using global signal regression (GSR) to remove spatially persistent artifact is controversial in neuroimaging (77, 78), but it remains a current field-wide standard (though see other recent and emerging approaches at (79, 80)). In order to address this controversy, we also include the mean effect and neural PCA results without GSR (**Fig. S18, Fig. S19**). In addition, when we compared the results of the neural PCA performed with and without GSR, we found that PC1 Δ GBC with GSR was significantly positively correlated with the PC2 Δ GBC without GSR (r=0.54, p<.001) (**Fig. S20E**). In contrast, PC1 Δ GBC with GSR was not significantly correlated with the PC1 Δ GBC without GSR (r=0.−0.04, p=0.28) (**Fig. S20D**). This indicates that when you perform a PCA on results without GSR, the first PC captures a substantial amount of the global signal that is removed when regressing out GSR.

### Neural Data Reduction via Functional Brain-wide Parcellation

All neural data analysed in this study is resting-state data. We examined the neural effects of ketamine at multiple levels of analysis: dense (91281 grayordinates); parcel (718 parcels); network (12 networks); and subcortex (5 subcortical structures). Psilocybin and LSD were analysed at the parcel-level only. Both the parcel-level and network-level data were acquired using a functionally-derived whole-brain parcellation via the recently-validated Cole-Anticevic Brain Network Parcellation (CAB-NP) atlas (81, 82). The subcortex-level data was acquired by parcellating the neural data using Freesurfer’s anatomically defined subcortical structures. All data was parcellated prior to running global brain connectivity, as this was shown to improve the signal-to-noise ratio (**Fig. S1**). While we report the mean effect of ketamine at grayordinate and parcel-level (**Fig. S21**), further analysis was conducted at the parcel-level as this gave the best trade-off between the sample size needed to resolve multivariate neurobehavioral solutions and the size of the feature space (52).

### Global Brain Connectivity

Global brain connectivity was calculated on the ketamine, psilocybin, and LSD datasets. Following preprocessing, the resting-state functional connectivity (FC) matrix was calculated for each participant by computing the Pearson’s correlation between every grayordinate in the brain with all other grayordinates. A Fisher’s r-to-Z transform was then applied. Global brain connectivity (GBC) was calculate by computing every grayordinate’s mean FC strength with all other grayordinates (i.e. the mean, per row, across all columns of the FC matrix). Thus, this calculation yielded a GBC map for each subject where each grayordinates value represents the mean connectivity of that grayordinate with all other grayordinates in the brain. GBC is a data-driven summary measure of connectedness that is unbiased with regards to the location of a possible alteration in connectivity (83) and is therefore a principled way for reducing the number of neural functional connectivity features while assessing neural variation across the entire brain. GBC was calculated as:

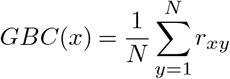

where *GBC*(*x*) denotes the GBC value at grayordinate *x*; *N* denotes the total number of grayordinates; 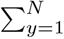 denotes the sum from *y* = 1 to *y* = *N* ; and where *r_xy_* denotes the correlation between the time-series of grayordinates *x* and *y*. For parcel-level and network-level maps, as outlined in the previous section we first computed the mean BOLD signal within each parcel/network for each participant, and then computed the pairwise FC between all parcels/networks. Finally, to obtain the parcellated GBC metric we computed the mean FC for each parcel/network.

### Principal Component Analysis of Neural Data

Global brain connectivity was calculated on the ketamine, psilocybin, and LSD datasets. For each dataset the input for the principal component analysis (PCA) of the neural data was the parcel-level Δ (substance - placebo) GBC maps for each subject. The PCA was computed using the 718 parcels across all participants. Significance of the neural PCA solution was assessed via permutation testing (1000 random shuffles of parcels within subject).

### Effective Dimensionality

Effective dimensionality was calculated on the ketamine, psilocybin, and LSD datasets to compare the dimensionality of the neural effects of different pharmacological substances. We used the participation ratio (PR), calculated as:

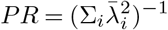

where {λ*_i_*} is the *ith* eigenvalue of the covariance matrix, and 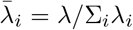 (54, 56). Larger values indicate a more complex higher dimensional dataset, while smaller values indicate a less complex lower dimensional dataset. Input for the calculation for the LSD condition were the Δ (LSD - placebo) GBC maps for each participant, input for the psilocybin condition were the Δ (psilocybin - placebo) GBC maps for each participant, and input for the ketamine condition were the Δ (ketamine - placebo) GBC maps for each participant. As sample size is relevant when calculating effective dimensionality, we first re-sampled the data ensuring sample size in each of the pharmacological conditions was always 22. We used either re-sampling or jackknifing to build a distribution of effective dimensionality values for each pharmacological condition. For LSD (N=24), we selected all 276 possible combinations of 22 participants and calculated effective dimensionality on each of the 276 subsamples to build a distribution. For psilocybin (N=23), used jackknifing to produce 22 subsamples of 22 participants and calculated effective dimensionality on each of the 22 subsamples to build a distribution. For ketamine (N=40) we randomly selected 100 subsamples from the 113380261800 possible combinations of 22 participants and calculated effective dimensionality on each of the 100 subsamples to build a distribution. To compare the pharmacological conditions we then ran a one-way ANOVA comparing effective dimensionality distributions across LSD, psilocybin, and ketamine. Finally we ran a series of post-hoc t-tests with Bonferroni correction.

### Neural Gene Expression Mapping

Neural gene expression mapping was calculated on the ketamine dataset only. Methods for the gene mapping analyses in this study are described in detail in prior publications (50, 52). To relate ketamine-specific n euroimaging e ffects t o t he c ortical t opography of gene expression for candidate receptors, we used cortical gene expression data from the publicly available Allen Human Brain Atlas (AHBA, RRID:SCR-007416), mapped to cortex (50). Specifically, t he A HBA q uantified expression levels across 20,737 genes obtained from six postmortem human brains using DNA microarray probes sampled from hundreds of neuroanatomical loci. We mapped gene expression on to 180 symmetrized cortical parcels from the HCP atlas (82) in line with recently published methods (52). This yielded a group-level map for each gene where the value in each parcel reflected the average expression level of that gene in the AHBA dataset. We selected two interneuron marker genes (somatostatin (SST) and parvalbumin (PVALB)) (84). As in prior works, we first excluded any gene expression maps where the cortical differential stability value was between 0 and +/− 0.1 (50). We then correlated the gene expression maps for each of the selected genes with our target map. As the gene expression maps are restricted to the cortex, the correlations were run on cortex only. To assess the significance of each correlation, we used the following approach, which is outlined in more detail in Burt et al. 2020 and **Figure S7**. We first generated 100,000 surrogate maps, whose spatial autocorrelation was matched to the spatial autocorrelation of the target map. The surrogate maps were generated separately for the left and right hemispheres and then merged together to produce 100,000 whole-brain cortical surrogate maps. For each selected gene, we correlated the gene expression map with each of the 100,000 surrogate maps to get a distribution of 100,000 simulated r values. We then used this distribution of simulated r values to calculate the significance of the correlation between the gene expression map and the target map. All p-values were FDR corrected.

### Principal Component Analysis of Behavioral Measures

The principal component analysis of behavioral measures was calculated on the ketamine dataset only. Subjective effects were analyzed using the Positive and Negative Syndrome Scale (PANSS), an assessment of psychosis symptom severity (85). The full PANSS battery is conventionally divided into three subscales: Positive symptom scale (7 items), Negative symptom scale (7 items), and General Psychopathology symptom scale (16 items). Cognition was analyzed using a spatial working memory paradigm, which resulted in a single cognition score. In total, this yielded 31 symptom variables per participant. Two participants had missing values for PANSS, so the mean was imposed. The principal component analysis (PCA) of behavioral data was computed using the 31 symptom variables across all N=40 participants. Variables were first scaled to have unit variance across participants before running the PCA. Significance of the derived principal components (PCs) was computed via permutation testing. For each permutation, participant order was randomly shuffled for each symptom variable before re-computing the PCA. This permutation was repeated 5,000 times to establish the null model. PCs which accounted for a proportion of variance that exceeded chance (p<0.05 across all 5000 permutations) were retained for further analysis.

### Mass Univariate Symptom-Neural Mapping

The mass univariate symptom-neural mapping was calculated on the ketamine dataset only. Methods for the mass univariate symptom-neural mapping are described in detail in prior publications (52). Behavioral scores were quantified in relation to individual GBC variation at the parcel-level via a mass univariate regression procedure. The resulting maps of regression coefficients reflects the strength of the relationship between participants’ behavioral score and Δ GBC at every neural location (718 parcels), across all 40 participants. The greater the magnitude of the coefficient for a given location, the stronger the statistical relationship between Δ GBC and the behavioral variation across participants. Significance of the maps was assessed via nonparametric permutation testing, 5000 random shuffles with TFCE (86) type-I error-protection computed via the Permutation Analysis of Linear Models program (87).

### Code Availability

Custom analysis codes written in Python, R, and Matlab are available from the corresponding author upon reasonable request.

