## Supplementary Materials for "Ketamine induces multiple individually distinct whole-brain functional connectivity signatures"

### Supplementary Tables

| Characteristic | Pre Ketamine (N=40) | | Post Ketamine (N=40) | | |
|---|---|---|---|---|---|
| | **Mean** | **S.D.** | **Mean** | **S.D.** | **P-Value** |
| **PANSS Positive Symptoms** | 7.51 | 0.84 | 14.21 | 3.50 | <.001** |
| **PANSS Negative Symptoms** | 7.95 | 1.03 | 14.57 | 5.49 | <.001** |
| **PANSS General Symptoms** | 17.14 | 1.81 | 29.92 | 6.98 | <.001** |
| **PANSS Total Psychopathology** | 32.59 | 2.66 | 58.72 | 14.27 | <.001** |
| **CADSS** | 0.10 | 0.38 | 16.78 | 10.32 | <.001** |
| **BDI** | 1.38 | 1.93 | 0.76 | 1.53 | 0.21 |

**Table S1. Retrospectively assessed (180 mins after drug administration) ketamine-induced subjective effects.** Effects were assessed using the Clinician Administered Dissociative States Scale (CADSS), the Positive and Negative Syndrome Scale for positive symptoms, negative symptoms and general psychopathology, and the Beck's Depression Inventory (BDI). N=40. ** indicates p<.001

DRAFT

| Characteristic | Healthy Participants (N=40) | |
|---|---|---|
| | **Mean** | **S.D.** |
| **Age (years)** | 25.73 | 3.74 |
| **Gender (% male)** | 0.73 | - |
| **Participant Education (years)** | 16.67 | 1.77 |
| **Maternal Education (years)** | 14.55 | 2.57 |
| **Paternal Education (years)** | 15.03 | 3.09 |
| **Smoking (% smokers)** | 0.03 | - |
| **Handedness (% right)** | 0.85 | - |
| **Race (%)** | Asian: 12.5% | |
| | Black: 22.5% | |
| | White: 55% | |
| | Mixed: 7.2% | |
| | Not Specified: 2.5% | |

**Table S2. Demographic Information.**

DRAFT

| | PC1 | PC2 |
|---|---|---|
| 1 Cognition - Spatial Working Memory | -0.002 | 0.131 |
| 2 PANSS P1 - Delusions | -0.157 | 0.330 |
| 3 PANSS P2 - Conceptual Disorganization | -0.267 | 0.024 |
| 4 PANSS P3 - Hallucinations | -0.022 | 0.335 |
| 5 PANSS P4 - Excitement | -0.163 | 0.109 |
| 6 PANSS P5 - Grandiosity | 0.023 | 0.262 |
| 7 PANSS P6 - Suspiciousness/Persecution | -0.069 | 0.033 |
| 8 PANSS P7 - Hostility | -0.150 | -0.072 |
| 9 PANSS N1 - Blunted Affect | -0.249 | -0.062 |
| 10 PANSS N2 - Emotional Withdrawal | -0.295 | -0.081 |
| 11 PANSS N3 - Poor Rapport | -0.086 | -0.302 |
| 12 PANSS N4 - Passive/Apathetic Social Withdrawal | -0.266 | -0.130 |
| 13 PANSS N5 - Difficulty in Abstract Thinking | -0.218 | 0.125 |
| 14 PANSS N6 - Lack of Spontaneity and Flow of Conversation | -0.277 | -0.152 |
| 15 PANSS N7 - Stereotyped Thinking | -0.159 | 0.129 |
| 16 PANSS G1 - Somatic Concern | -0.034 | 0.147 |
| 17 PANSS G2 - Anxiety | -0.167 | 0.023 |
| 18 PANSS G3 - Guilt Feelings | -0.025 | 0.035 |
| 19 PANSS G4 - Tension | -0.197 | -0.044 |
| 20 PANSS G5 - Mannerisms and Posturing | -0.132 | 0.040 |
| 21 PANSS G6 - Depression | -0.137 | 0.140 |
| 22 PANSS G7 - Motor Retardation | -0.236 | -0.040 |
| 23 PANSS G8 - Uncooperativeness | -0.075 | -0.369 |
| 24 PANSS G9 - Unusual Thought Content | -0.126 | 0.305 |
| 25 PANSS G10 - Disorientation | -0.202 | 0.144 |
| 26 PANSS G11 - Poor Attention | -0.245 | -0.053 |
| 27 PANSS G12 - Lack of Judgement and Insight | -0.256 | 0.043 |
| 28 PANSS G13 - Disturbance of Volition | -0.197 | -0.032 |
| 29 PANSS G14 - Poor Impulse Control | -0.059 | -0.362 |
| 30 PANSS G15 - Preoccupation | -0.142 | 0.219 |
| 31 PANSS G16 - Active Social Avoidance | -0.250 | -0.132 |

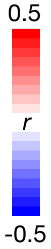


**Table S3. Delta Behavioral Principal Component (PCA) Loadings.** Table of the loadings of each of the 31 behavioral items on the two significant PCs (also seen in radar plot in **Fig. 3D**). Green = cognition, purple = PANSS positive, blue = PANSS negative, pink = PANSS general. Positive loadings are indicated in red; negative loadings are shown in blue.

### Supplementary Figures

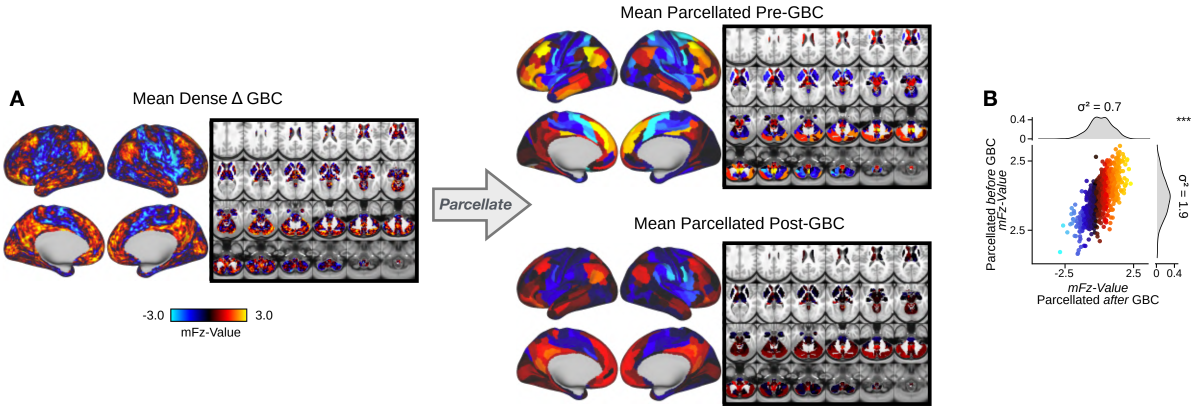


**Fig. S1. Parcellating before running GBC results in an improved signal to noise ratio. (A)** (Left) Unthresholded $\Delta$ (ketamine - placebo) GBC mean map at the grayordinate-level. (Right) Unthresholded $\Delta$ GBC mean map parcellated before running GBC (top) or after (bottom) running GBC. **(B)** Scatterplot showing relationship between parcel values (GBC mFz-values) computed pre vs. post GBC. Marginal distribution plots show the variance of $\Delta$ GBC mean map parcellated before GBC ($\sigma^2$ = 1.9, right) and the variance of $\Delta$ GBC mean map parcellated before GBC ($\sigma^2$ = 0.7, top). A paired Pitman-Morgan test revealed that there is a significant difference in variance between the two maps (p<0.001).

DRAFT

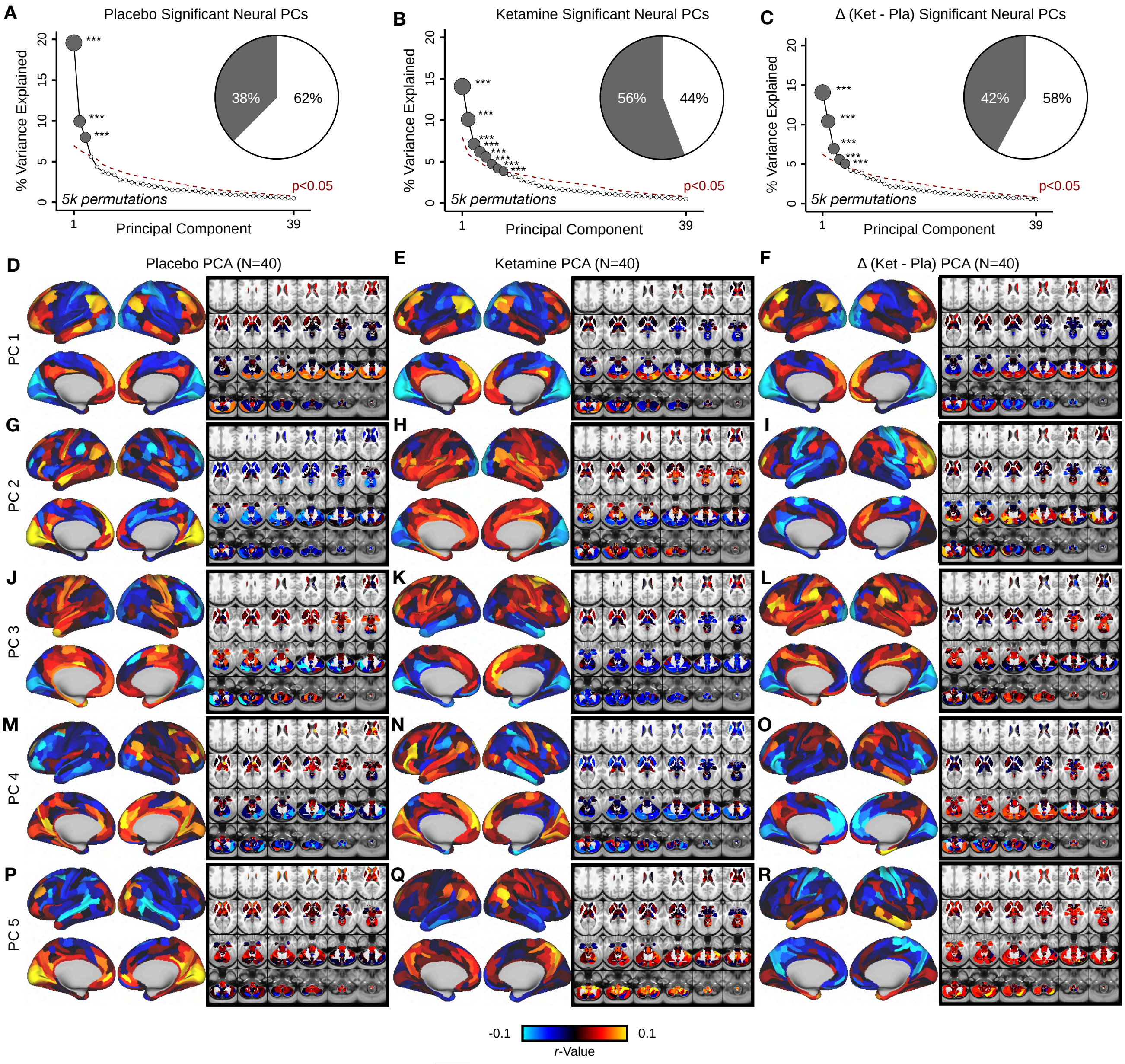


**Fig. S2. Principal component analysis (PCA) on the neural features from the placebo, ketamine, and △ (ketamine - placebo) neural GBC maps. (A)** Screeplot showing the total proportion of variance explained by the 39 PCs from the PCA performed across all 718 neural features in 40 subjects in the placebo condition. The size of each gray point is proportional to the variance explained by that PC. The first 3 gray PCs were determined to be significant using a permutation test (5000 permutations). Inset shows the proportion of variance both accounted and not accounted for by the 3 PCs. Together, these 3 PCs captures 37.5% of the total variation in neural GBC in the sample. **(B)** Screeplot showing the total proportion of variance explained by the 39 PCs from the PCA performed across all 718 neural features in 40 subjects in the ketamine condition. The size of each gray point is proportional to the variance explained by that PC. The first 8 gray PCs were determined to be significant using a permutation test 5000 permutations). Inset shows the proportion of variance both accounted and not accounted for by the 8 PCs. Together, these 8 PCs captures 55.8% of the total variation in neural GBC in the sample. **(C)** Screeplot showing the total proportion of variance explained by the 39 PCs from the PCA performed across all 718 neural features in 40 subjects in the △ condition. The size of each gray point is proportional to the variance explained by that PC. The first 5 gray PCs were determined to be significant using a permutation test (5000 permutations). Inset shows the proportion of variance both accounted and not accounted for by the 5 PCs. Together, these 5 PCs captures 42.1% of the total variation in neural GBC in the sample. **(D-R)** Neural maps for PCs 1-5 from the PCA performed on neural features (718 whole-brain parcel GBC) across all subjects in the placebo (left column), ketamine (centre column), and △ (ketamine - placebo) conditions (right column). Red/orange areas indicate parcels that have a high positive loading score onto PC, while blue areas indicate parcels that have a high negative loading score onto PC.

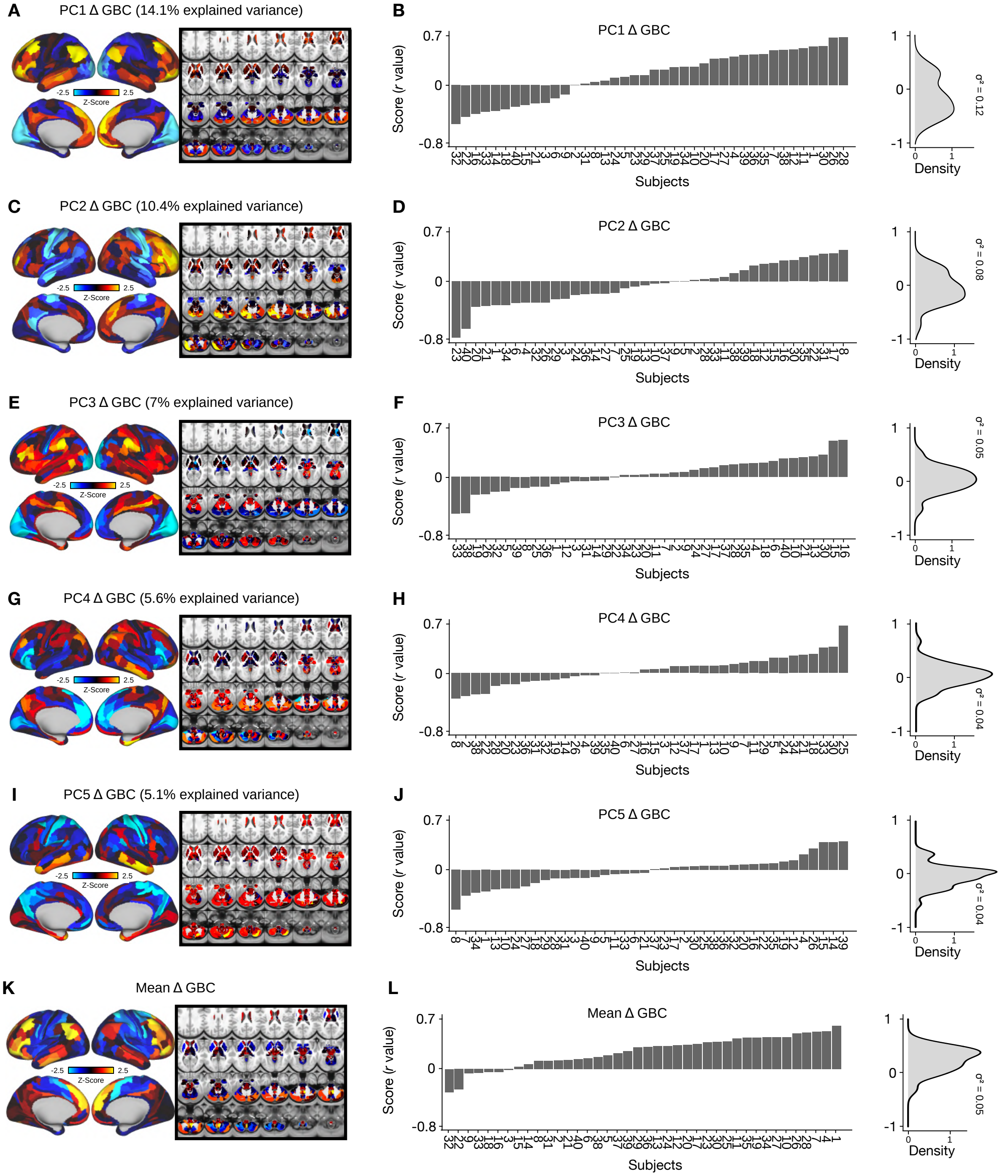


**Fig. S3. Individual variation in PC1-5 △ GBC and mean △ GBC. (A, C, E, G, I)** Unthresholded PC1-5 △ GBC Z-score map (Z-scores computed across 718 parcels). Red/orange areas indicate parcels that have a high positive loading score onto PC1, while blue areas indicate parcels that have a high negative loading score onto PC1. (PC1 and PC2 are sign flipped for visual comparison with mean). **(B, D, F, H, J)** (Left) Bar plot showing the PC1-5 △ GBC score (r-value) for each participant (N=40). The score for each participant was calculated by correlating their individual △ GBC map with the PC △ GBC map. (Right) density plot showing the variation captured by each PC △ GBC. **(K)** Unthresholded mean △ (ketamine - placebo) GBC Z-score map (Z-scores computed across 718 parcels). Red/orange areas indicate regions where participants exhibited stronger GBC in the ketamine condition, whereas blue areas indicate regions where participants exhibited reduced GBC in the ketamine condition, compared with the placebo condition. **(L)** (Left) Bar plot showing the mean △ GBC score (r-value) for each participant (N=40). The score for each participant was calculated by correlating their individual △ GBC map with the mean △ GBC map. (Right) density plot showing the variation captured by the mean △ GBC.

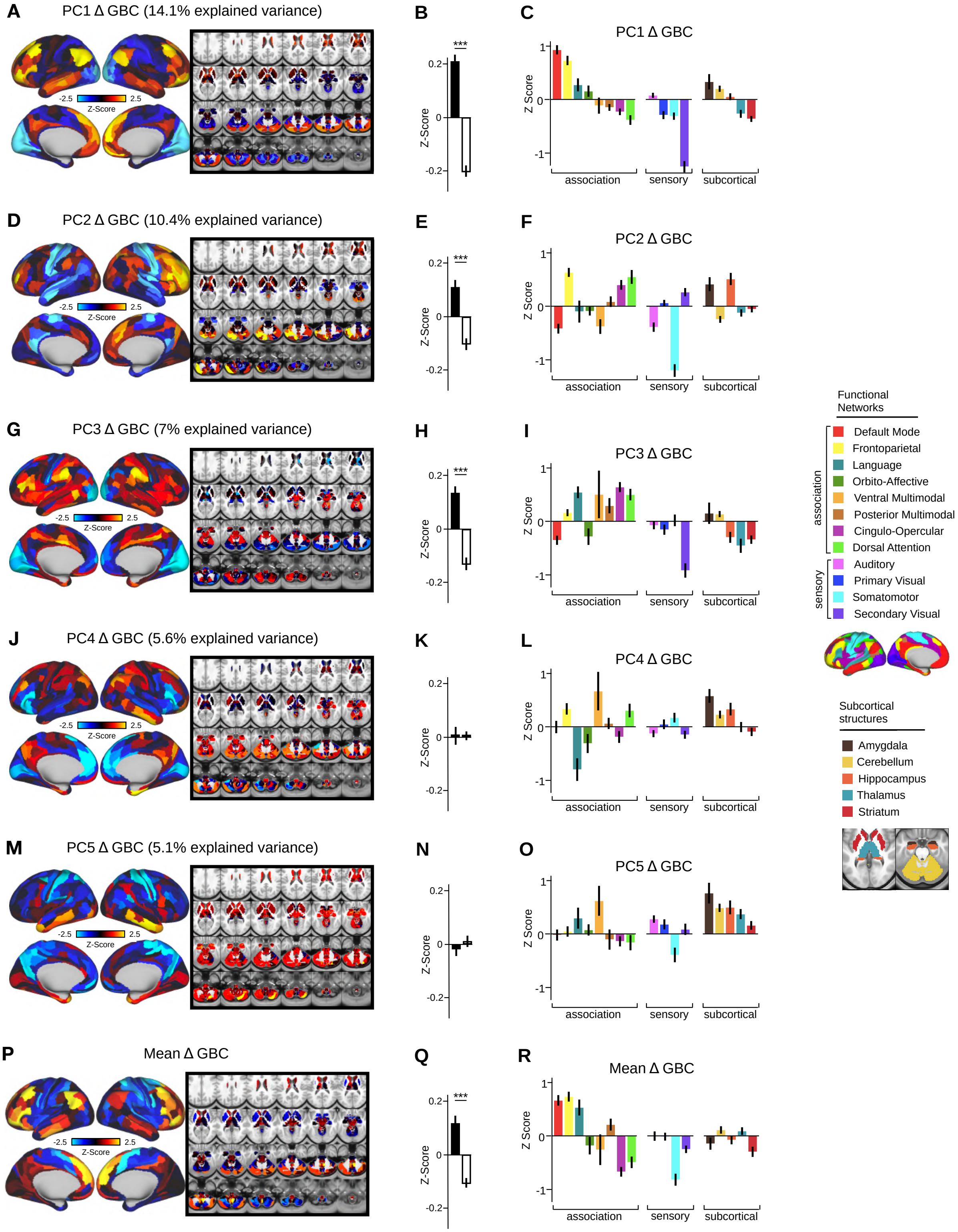


**Fig. S4. Network breakdown of PC1-5 △ GBC and mean △ GBC. (A,D,G,J,M)** Unthresholded PC1-5 △ GBC Z-score map (Z-scores computed across 718 parcels). Red/orange areas indicate parcels that have a high positive loading score onto the PC, while blue areas indicate parcels that have a high negative loading score onto the PC. (PC1 and PC2 are sign flipped for visual comparison with mean). **(B,E,H,K,N)** Bar plot showing the PC1-5 (△ GBC Z-score across association (black) and sensory (white) network parcels. See graphs right for association and sensory network groupings. **(C,F,I,L,O)** Bar plot showing the mean Z-score for each network and each anatomical subcortical structure for PC1-5 △ GBC (see inset right for color labels). Networks are grouped into association and sensory networks. **(P)** Unthresholded mean △ (ketamine - placebo) GBC Z-score map (Z-scores computed across 718 parcels). Red/orange areas indicate regions where participants exhibited stronger GBC in the ketamine condition, whereas blue areas indicate regions where participants exhibited reduced GBC in the ketamine condition, compared with the placebo condition. **(Q)** Bar plot showing the mean (△ GBC Z-score across association (black) and sensory (white) network parcels. See graphs right for association and sensory network groupings. **(R)** Bar plot showing the mean Z-score for each network and each anatomical subcortical structure for mean △ GBC (see inset right for color labels). Networks are grouped into association and sensory networks.

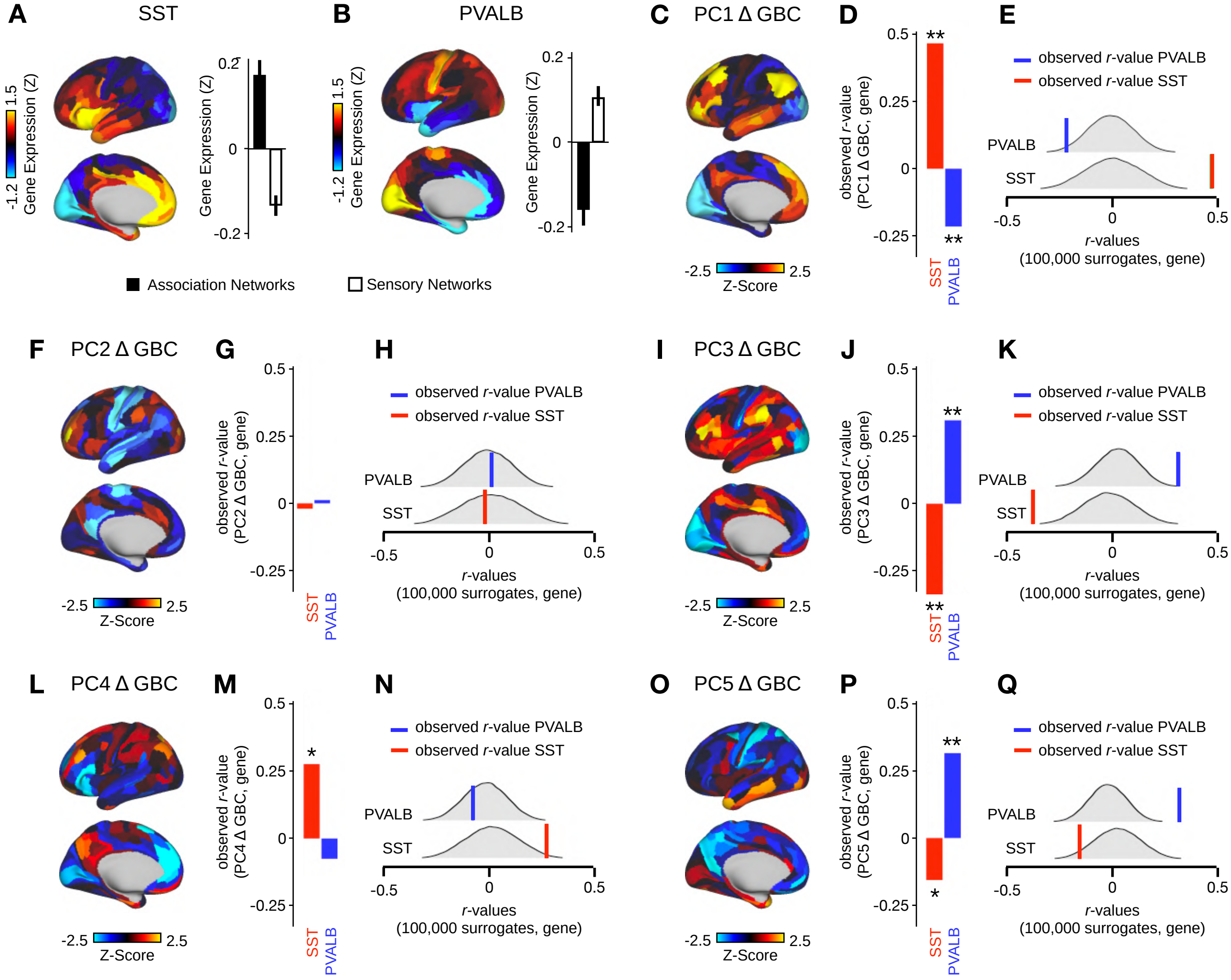


**Fig. S5. PC1 △ GBC and PC3-5 △ GBC maps tracks SST and PVALB neural gene expression patterns. (A)** Gene expression pattern of interneuron marker gene somatostatin (SST). Left: positive (yellow) regions show areas where the gene of interest is highly expressed, whereas negative (blue) regions indicate low expression values. Right: bar plot showing the mean gene expression Z-score across association (black) and sensory (white) networks. **(B)** Gene expression pattern of interneuron marker parvalbumin (PVALB). Left: positive (yellow) regions show areas where the gene of interest is highly expressed, whereas negative (blue) regions indicate low expression values. Right: bar plot showing the mean gene expression Z-score across association (black) and sensory (white) networks. **(C,F,I,L,O)** Unthresholded PC1-5 △ GBC Z-score map (Z-scores computed across 718 parcels). Red/orange areas indicate parcels that have a high positive loading score onto the PC, while blue areas indicate parcels that have a high negative loading score onto the PC. (PC1 and PC2 are sign flipped for visual comparison with mean). **(D,G,J,M,P)** Bar plots showing the correlation between PC1-5 △ GBC and the following gene expression maps: SST (red) and PVALB (blue). All p-values are FDR corrected. ** = p<.001, * = p<.05. **(E,H,K,N,Q)** Distribution of 100,000 simulated r values for SST (bottom) and PVALB (top). Bold lines indicate the observed r value between the PC △ GBC and SST (red) and PVALB (blue).

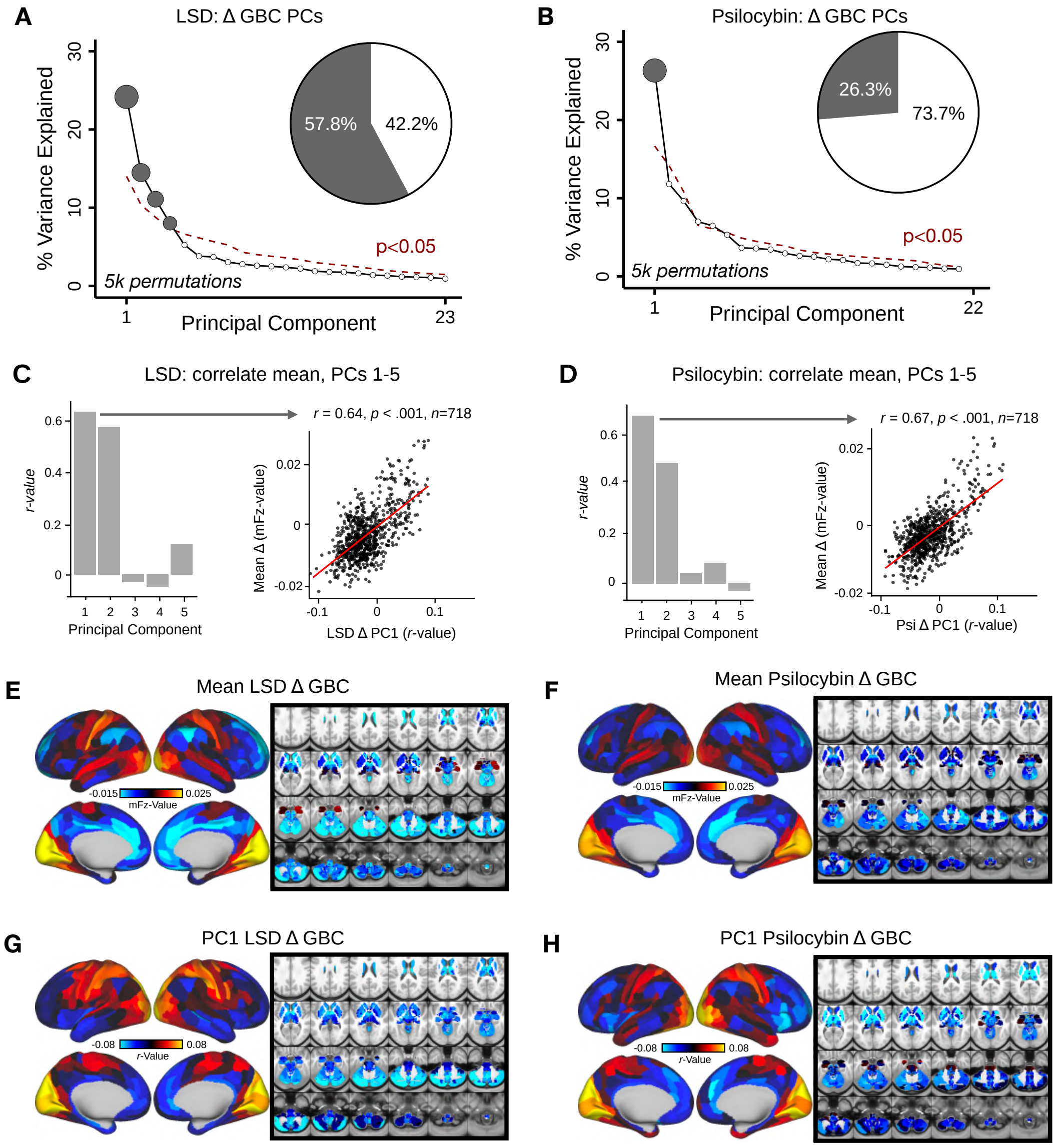


**Fig. S6. Principal Component Analysis of LSD and psilocybin △ GBC (A)** Results of PCA performed on △ (LSD - placebo) GBC neural features (718 whole-brain parcel GBC) across all subjects (N=24). Screeplot showing the % variance explained by the each of the 23 PCs. The first 4 PCs (dark grey) were determined to be significant using a permutation test (p<.05, 5000 permutations). The size of each dark grey point is proportionate to the variance explained. Inset shows the proportion of variance both accounted and not accounted for by the 4 PCs. Together, these 3 PCs captures 57.8% of the total variation in neural GBC in the sample. **(B)** Results of PCA performed on △ (psilocybin - placebo) GBC neural features (718 whole-brain parcel GBC) across all subjects (N=23). Screeplot showing the % variance explained by the each of the 22 PCs. The first PC (dark grey) was determined to be significant using a permutation test (p<.05, 5000 permutations). The size of the dark grey point is proportionate to the variance explained. Inset shows the proportion of variance both accounted and not accounted for by the PC. The single PC captures 26.3% of the total variation in neural GBC in the sample. **(C)** Bar plot showing the correlation between each of the first five LSD △ GBC PCs and the mean LSD △ GBC. r values for PC1 and PC2 are significant (p<.001, Bonferroni corrected). (Inset) Scatterplot showing the relationship across parcels between mean LSD △ GBC and PC1 LSD △ GBC maps. **(D)** Bar plot showing the correlation between each of the first five psilocybin △ GBC PCs and the mean psilocybin △ GBC map. r values for PC1 and PC2 are significant (p<.001, Bonferroni corrected). (Inset) Scatterplot showing the relationship across parcels between mean psilocybin △ GBC and PC1 psilocybin △ GBC maps. **(E)** Unthresholded mean LSD △ GBC Z-score map at the parcel level (No. parcels = 718) (Z-scores computed across 718 parcels). Red/orange areas indicate regions where participants exhibited stronger GBC in the LSD condition, whereas blue areas indicate regions where participants exhibited reduced GBC in the LSD condition, compared with the placebo condition. **(F)** Unthresholded mean psilocybin △ GBC Z-score map at the parcel level (No. parcels = 718) (Z-scores computed across 718 parcels). Red/orange areas indicate regions where participants exhibited stronger GBC in the psilocybin condition, whereas blue areas indicate regions where participants exhibited reduced GBC in the psilocybin condition, compared with the placebo condition. **(H)** PC1 LSD △ GBC Z-score map (Z-scores computed across 718 parcels). PC1 LSD △ GBC explains 25% of all variance. Red/orange areas indicate parcels that have a high positive loading score onto PC1, while blue areas indicate parcels that have a high negative loading score onto PC1. (PC1 is sign flipped for visual comparison with mean). **(I)** PC1 psilocybin △ GBC Z-score map (Z-scores computed across 718 parcels). PC1 △ GBC explains 26.3% of all variance. Red/orange areas indicate parcels that have a high positive loading score onto PC1, while blue areas indicate parcels that have a high negative loading score onto PC1.

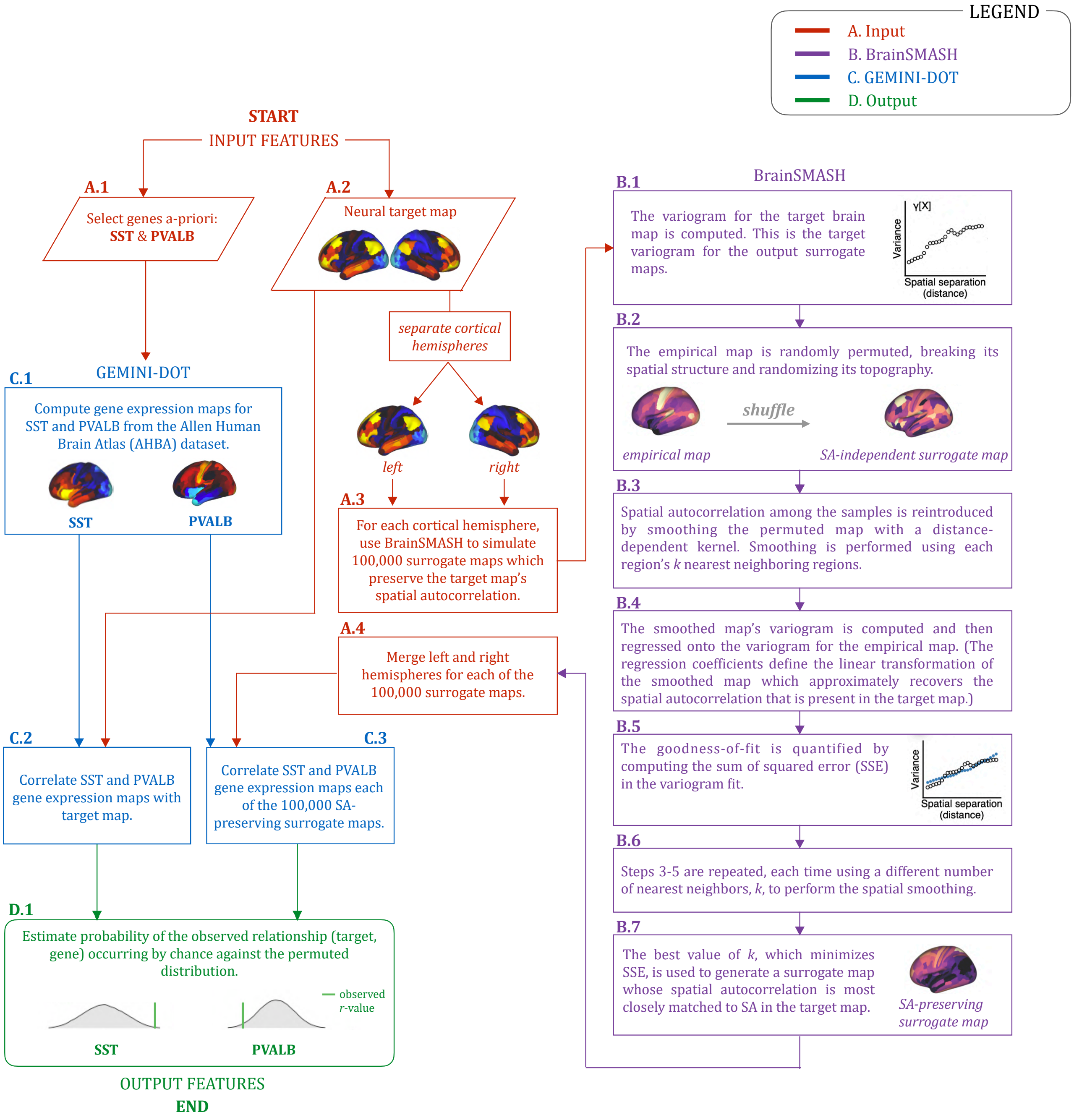


**Fig. S7. Gene analysis workflow.** Gene analysis workflow outlining: (A.1-2) the input features (red); (B.1-7) the generation of surrogate maps using BrainSMASH (purple); (C.1-3) the generation of gene expression maps using GEMINI-DOT (blue); and (D.1) the ouput features (green).

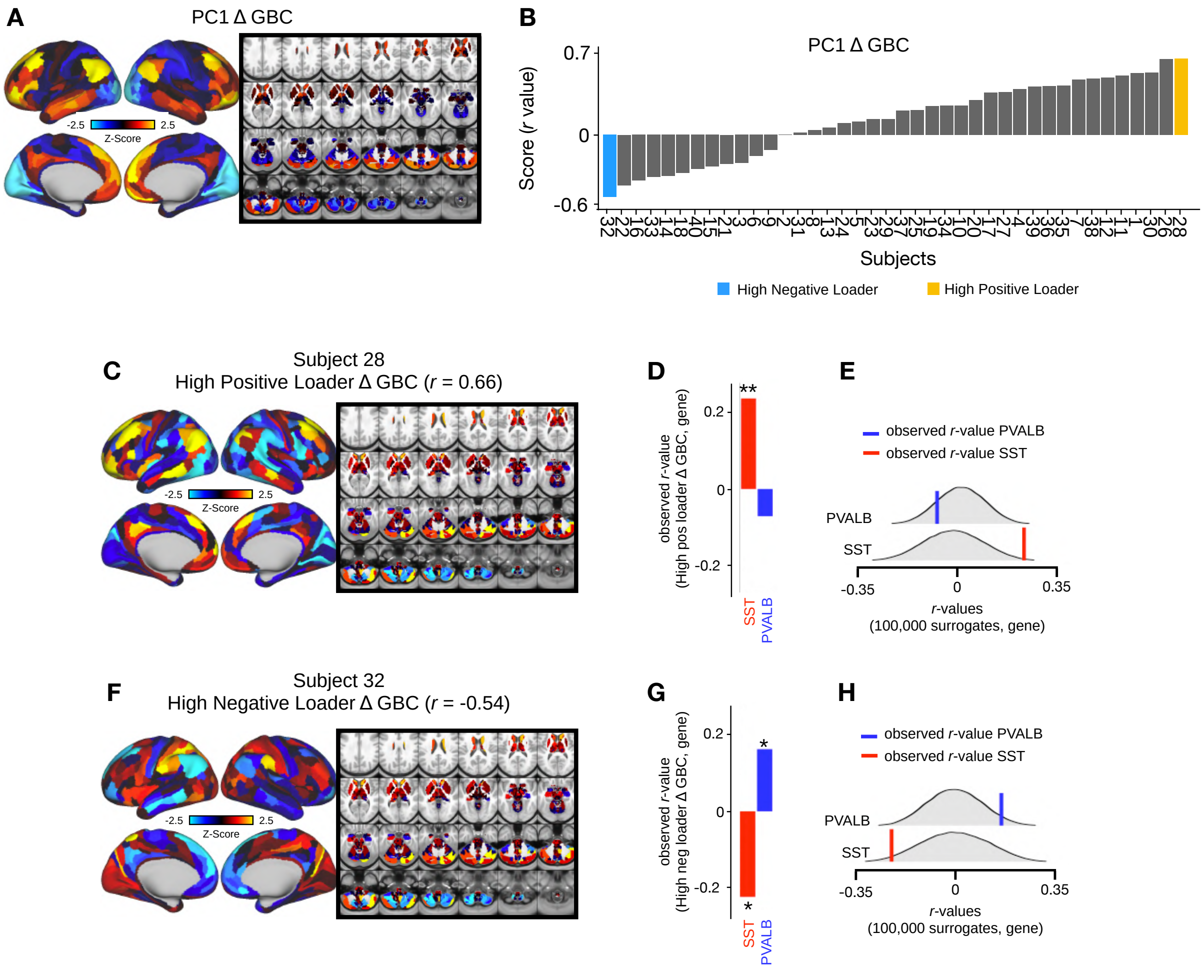


**Fig. S8. High positive and negative PC1 △ GBC loaders track SST and PVALB gene expression topographies. (A)** Unthresholded PC1 △ GBC Z-score map (Z-scores computed across 718 parcels). PC1 △ GBC explains 14.1% of all variance. Red/orange areas indicate parcels that have a high positive loading score onto PC1, while blue areas indicate parcels that have a high negative loading score onto PC1. (PC1 is sign flipped for visual comparison with mean). **(B)** Bar plot showing the PC1 △ GBC score (r-value) for each participant (N=40). For each participant the the PC1 △ GBC score is calculated by correlating their individual △ GBC map with the PC1 △ GBC map. Blue = high negative PC1 loader, yellow = high positive PC1 loader. **(C)** △ GBC map for the high positive loader (subject 28). Red/orange areas indicate parcels in which there is a increased GBC following ketamine, while blue areas indicate parcels in which there is decreased GBC following ketamine. **(D)** Bar plot showing the correlation between subject 28 △ GBC and the following gene expression maps: SST (r=0.23, p=.009) (red) and PVALB (r=-0.07, p=.189) (blue). All p-values are FDR corrected. **(E)** Distribution of 100,000 simulated r values for subject 28 △ GBC and SST (bottom) and subject 28 △ GBC and PVALB (top). Bold lines indicate the observed r value between PC1 △ GBC and SST (red) and PVALB (blue). **(F)** △ GBC map for the high negative loader (subject 32). Red/orange areas indicate parcels in which there is a increased GBC following ketamine, while blue areas indicate parcels in which there is decreased GBC following ketamine. **(G)** Bar plot showing the correlation between subject 32 △ GBC and the following gene expression maps: SST (r=-0.22, p=.046) (red) and PVALB (r=0.16, p=.056) (blue). All p-values are FDR corrected. **(H)** Distribution of 100,000 simulated r values for subject 32 △ GBC and SST (bottom) and subject 32 △ GBC and PVALB (top). Bold lines indicate the observed r value between PC1 △ GBC and SST (red) and PVALB (blue).

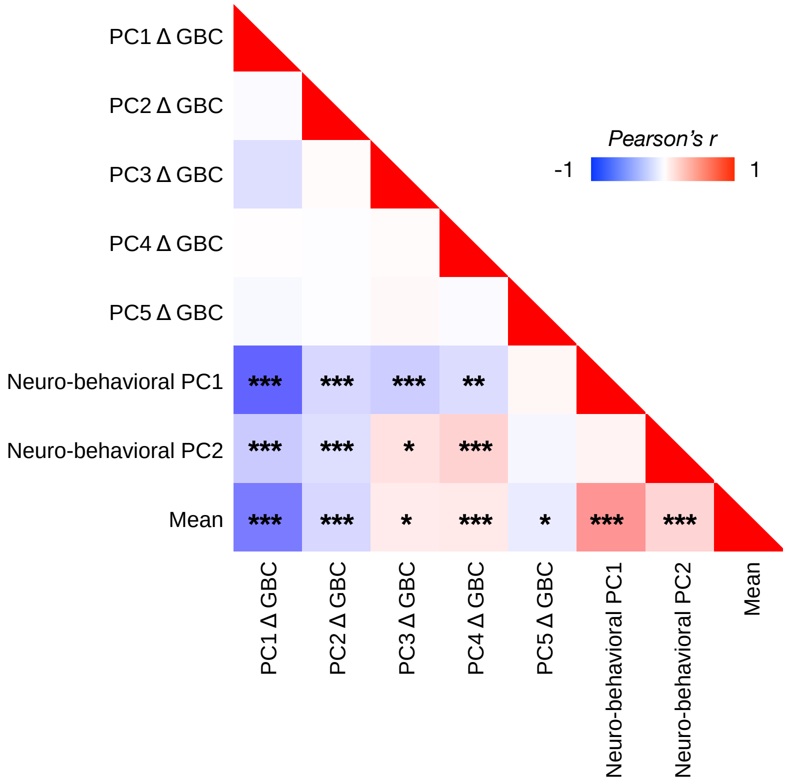


**Fig. S9. Correlations between PC1-5 △ GBC, neuro-behavioral PC1-2, and mean △ GBC maps.** The PC1 △ GBC map is highly negatively correlated with the neuro-behavioral PC1 map (r=-0.62) and the mean △ GBC map (r=-0.56), while neural-behavioral PC2 map is moderately positively correlated with the mean △ GBC map (r=0.41). * = p<.05, ** = p<.01, *** = p<.001.

DRAFT

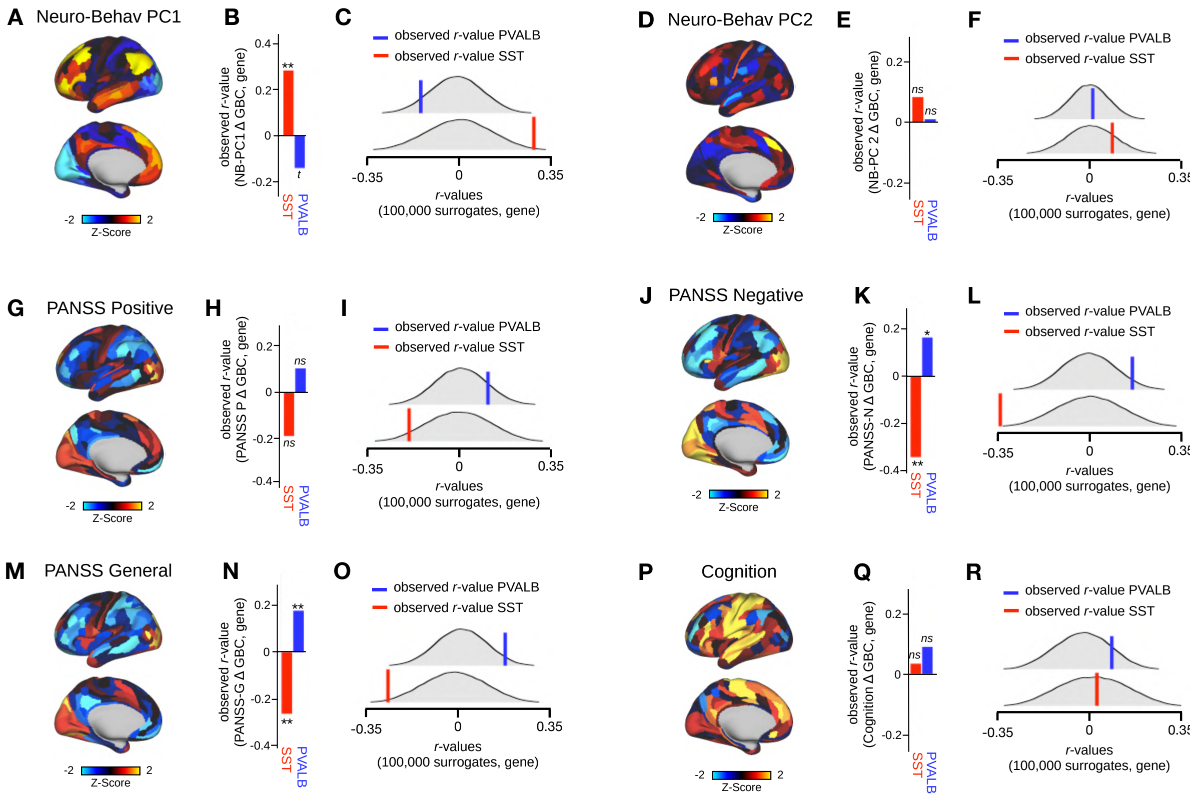


**Fig. S10. Neuro-behavioral PC1, PANSS negative, and PANSS general track SST and PVALB gene expression topographies.** **(A)** Neuro-behavioral PC1 map showing the relationship between the behavioral PC1 score for each participant regressed onto the $\Delta$ GBC map for each participant (N=40). **(B)** Bar plot showing the correlation between the neuro-behavioral PC1 map and the following gene expression maps: SST (r=0.29, p=.009) (red) and PVALB (r=-0.14, p=.089) (blue). All p-values are FDR corrected. **(C)** Distribution of 100,000 simulated r values for neuro-behavioral PC1 and: SST (bottom), PVALB (top). Bold lines indicate the observed r value between neuro-behavioral PC1 and: SST (red), PVALB (blue). **(D)** Neuro-behavioral PC2 map showing the relationship between the behavioral PC2 score for each participant regressed onto the $\Delta$ GBC map for each participant. **(E)** Bar plot showing the correlation between the neuro-behavioral PC2 map and the following gene expression maps: SST (r=0.09, p=.22) (red) and PVALB (r=0.01, p=.75) (blue). All p-values are FDR corrected. **(F)** Distribution of 100,000 simulated r values for neuro-behavioral PC2 and: SST (bottom), PVALB (top). Bold lines indicate the observed r value between PANSS negative and: SST (red), PVALB (blue). **(G)** PANSS positive map showing the relationship between the PANSS positive score for each participant regressed onto the $\Delta$ GBC map for each participant. **(H)** Bar plot showing the correlation between the PANSS positive map and the following gene expression maps: SST (r=-0.19, p=.103) (red) and PVALB (r=0.12, p=.164) (blue). All p-values are FDR corrected. **(I)** Distribution of 100,000 simulated r values for PANSS positive and: SST (bottom), PVALB (top). Bold lines indicate the observed r value between PANSS positive and: SST (red), PVALB (blue). **(J)** PANSS negative map showing the relationship between the PANSS negative score for each participant regressed onto the $\Delta$ GBC map for each participant. **(K)** Bar plot showing the correlation between the PANSS negative map and the following gene expression maps: SST (r=-0.34, p<.001) (red) and PVALB (r=0.16, p=.05) (blue). All p-values are FDR corrected. **(L)** Distribution of 100,000 simulated r values for PANSS negative and: SST (bottom), PVALB (top). Bold lines indicate the observed r value between PANSS negative and: SST (red), PVALB (blue). **(M)** PANSS general map showing the relationship between the PANSS general score for each participant regressed onto the $\Delta$ GBC map for each participant. **(N)** Bar plot showing the correlation between the PANSS general map and the following gene expression maps: SST (r=-0.26, p=.019) (red) and PVALB (r=0.18, p=.04) (blue). All p-values are FDR corrected. **(O)** Distribution of 100,000 simulated r values for PANSS general and: SST (bottom), PVALB (top). Bold lines indicate the observed r value between PANSS general and: SST (red), PVALB (blue). **(P)** Cognition map showing the relationship between the cognition score for each participant regressed onto the $\Delta$ GBC map for each participant. **(Q)** Bar plot showing the correlation between the cognition map and the following gene expression maps: SST (r=0.03, p=.84) (red) and PVALB (r=0.09, p=.33) (blue). All p-values are FDR corrected. **(R)** Distribution of 100,000 simulated r values for cognition and: SST (bottom), PVALB (top). Bold lines indicate the observed r value between cognition and: SST (red), PVALB (blue).

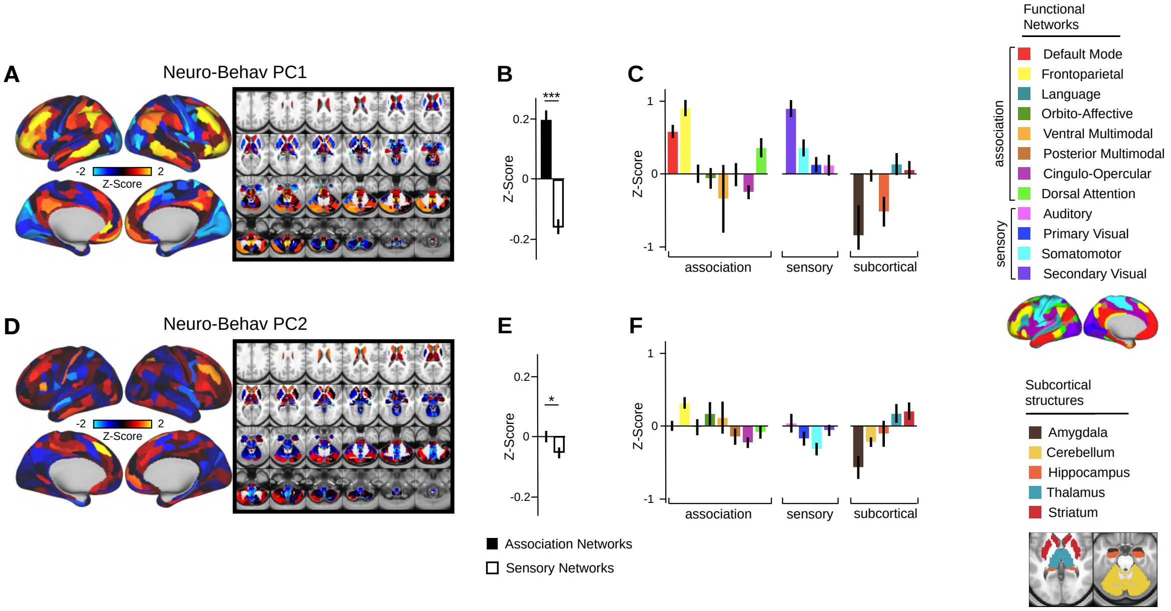


**Fig. S11. Network breakdown of neuro-behavioral PC1 and neuro-behavioral PC2. (A)** Neuro-behavioral PC1 map showing the relationship between the behavioral PC1 score for each participant regressed onto the $\Delta$ GBC map for each participant (N=40). Values shown in each brain parcel are the Z-scored regression coefficient (behavioral PC1 score, $\Delta$ GBC) across all 40 subjects. Red/orange areas indicate parcels in which there is a positive relationship between GBC and the behavioral PC1 score, while blue areas indicate parcels in which there is a negative relationship between GBC and the behavioral PC1 score. **B)** Bar plot showing the neuro-behavioral PC1 GBC Z-score across association (black) and sensory (white) network parcels. **(C)** Bar plot showing the mean correlation ($\Delta$ GBC, behavioral PC1 score) for each network and each anatomical subcortical structure (see inset right for color labels). **(D)** Neuro-behavioral PC2 map showing the relationship between the behavioral PC2 score for each participant regressed onto the $\Delta$ GBC map for each participant (N=40). Values shown in each brain parcel are the Z-scored regression coefficient (behavioral PC2 score, $\Delta$ GBC) across all 40 subjects. Red/orange areas indicate parcels in which there is a positive relationship between GBC and the behavioral PC2 score, while blue areas indicate parcels in which there is a negative relationship between GBC and the behavioral PC2 score. **(E)** Bar plot showing the neuro-behavioral PC2 GBC Z-score across association (black) and sensory (white) network parcels. **(F)** Bar plot showing the mean correlation ($\Delta$ GBC, behavioral PC2 score) for each network and each anatomical subcortical structure (see inset right for color labels).

DRAFT

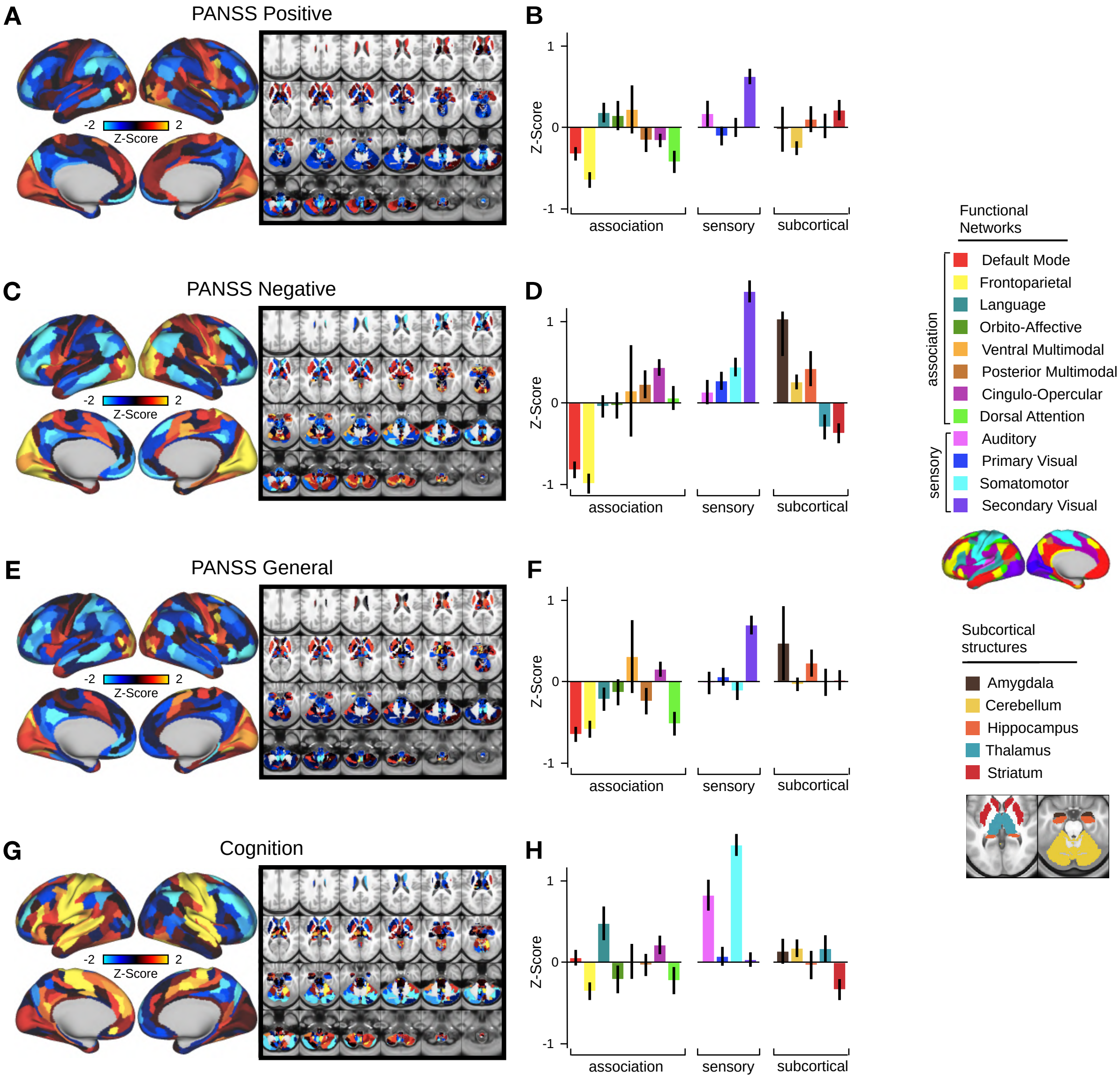


**Fig. S12. Network breakdown of existing behavioral measures: 3-factor PANSS subscales and cognition Δ GBC maps. (A)** PANSS positive map showing the relationship between the PANSS positive score for each participant regressed onto the Δ GBC map for each participant (N=40). Values shown in each brain parcel are the Z-scored regression coefficient (PANSS positive, Δ GBC) across all 40 subjects. Red/orange areas indicate parcels in which there is a positive relationship between GBC and the PANSS positive score, while blue areas indicate parcels in which there is a negative relationship between GBC and the PANSS positive score. **(B)** Bar plot showing the mean correlation (Δ GBC, PANSS positive score) for each network and each anatomical subcortical structure (see inset right for color labels). **(C)** PANSS negative map showing the relationship between the PANSS negative score for each participant regressed onto the Δ GBC map for each participant (N=40). Values shown in each brain parcel are the Z-scored regression coefficient (PANSS negative, Δ GBC) across all 40 subjects. Red/orange areas indicate parcels in which there is a positive relationship between GBC and the PANSS negative score, while blue areas indicate parcels in which there is a negative relationship between GBC and the PANSS negative score. **(D)** Bar plot showing the mean correlation (Δ GBC, PANSS negative score) for each network and each anatomical subcortical structure (see inset right for color labels). **(E)** PANSS general map showing the relationship between the PANSS general score for each participant regressed onto the Δ GBC map for each participant (N=40). Values shown in each brain parcel are the Z-scored regression coefficient (PANSS general, Δ GBC) across all 40 subjects. Red/orange areas indicate parcels in which there is a positive relationship between GBC and the PANSS general score, while blue areas indicate parcels in which there is a negative relationship between GBC and the PANSS general score. **(F)** Bar plot showing the mean correlation (Δ GBC, PANSS general score) for each network and each anatomical subcortical structure (see inset right for color labels). **(G)** Cognition map showing the relationship between the cognition score for each participant regressed onto the Δ GBC map for each participant (N=40). Values shown in each brain parcel are the Z-scored regression coefficient (cognition, Δ GBC) across all 40 subjects. Red/orange areas indicate parcels in which there is a general relationship between GBC and the cognition score, while blue areas indicate parcels in which there is a negative relationship between GBC and the cognition score. **(L)** Bar plot showing the mean correlation (Δ GBC, cognition score) for each network and each anatomical subcortical structure (see inset right for color labels).

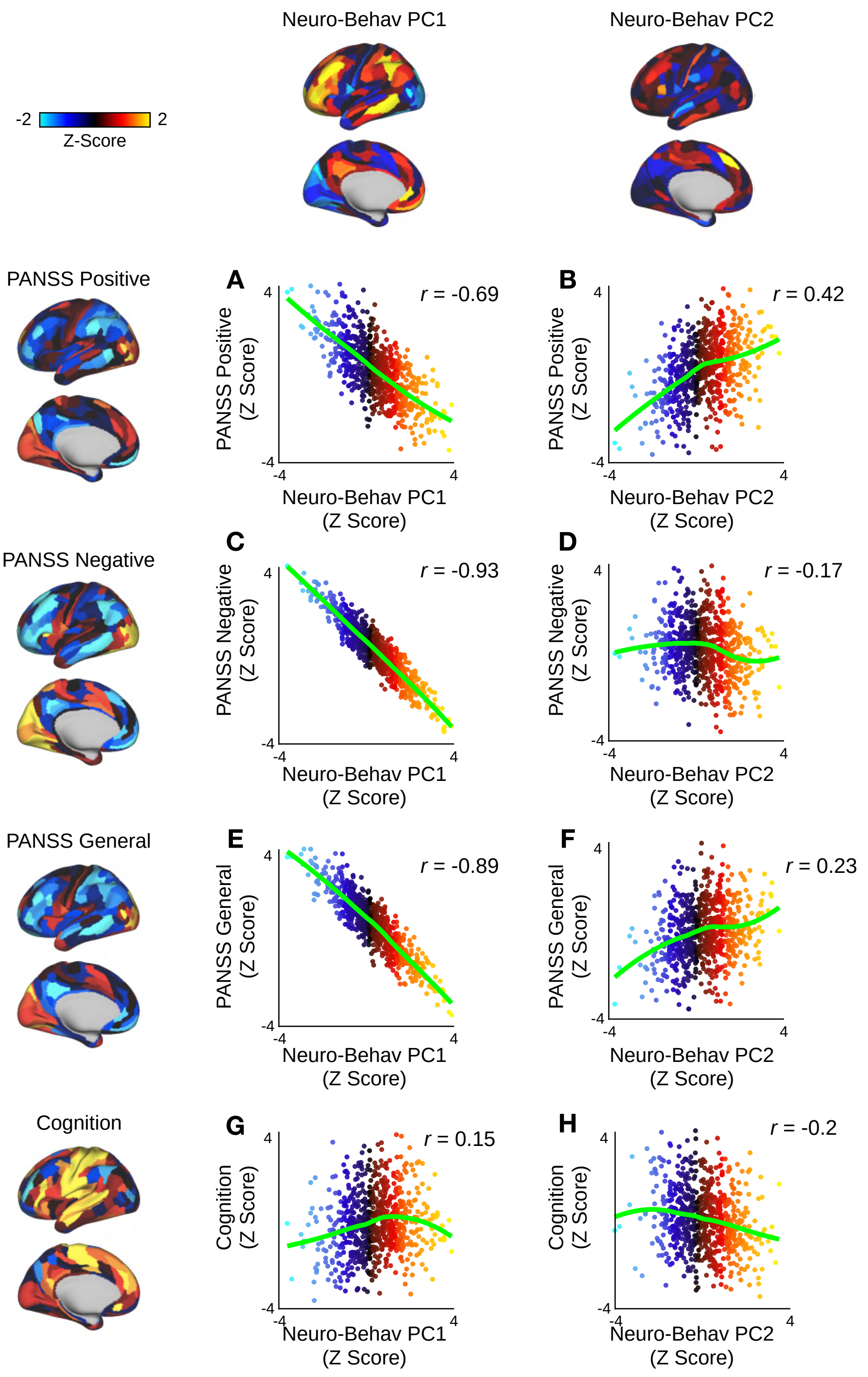


**Fig. S13. Scatter plots showing the relationship between neuro-behavioral PC1-2 and the 3-factor PANSS subscales. (A)** Scatter plot showing the relationship between PANSS positive (left) and neuro-behavioral PC1 (top). **(B)** Scatter plot showing the relationship between PANSS positive (left) and neuro-behavioral PC2 (top). **(C)** Scatter plot showing the relationship between PANSS negative (left) and neuro-behavioral PC1 (top). **(D)** Scatter plot showing the relationship between PANSS negative (left) and neuro-behavioral PC2 (top). **(E)** Scatter plot showing the relationship between PANSS general (left) and neuro-behavioral PC1 (top). **(F)** Scatter plot showing the relationship between PANSS general (left) and neuro-behavioral PC2 (top). **(G)** Scatter plot showing the relationship between cognition (left) and neuro-behavioral PC1 (top) (r=0.15, p . **(H)** Scatter plot showing the relationship between cognition (left) and neuro-behavioral PC2 (top).

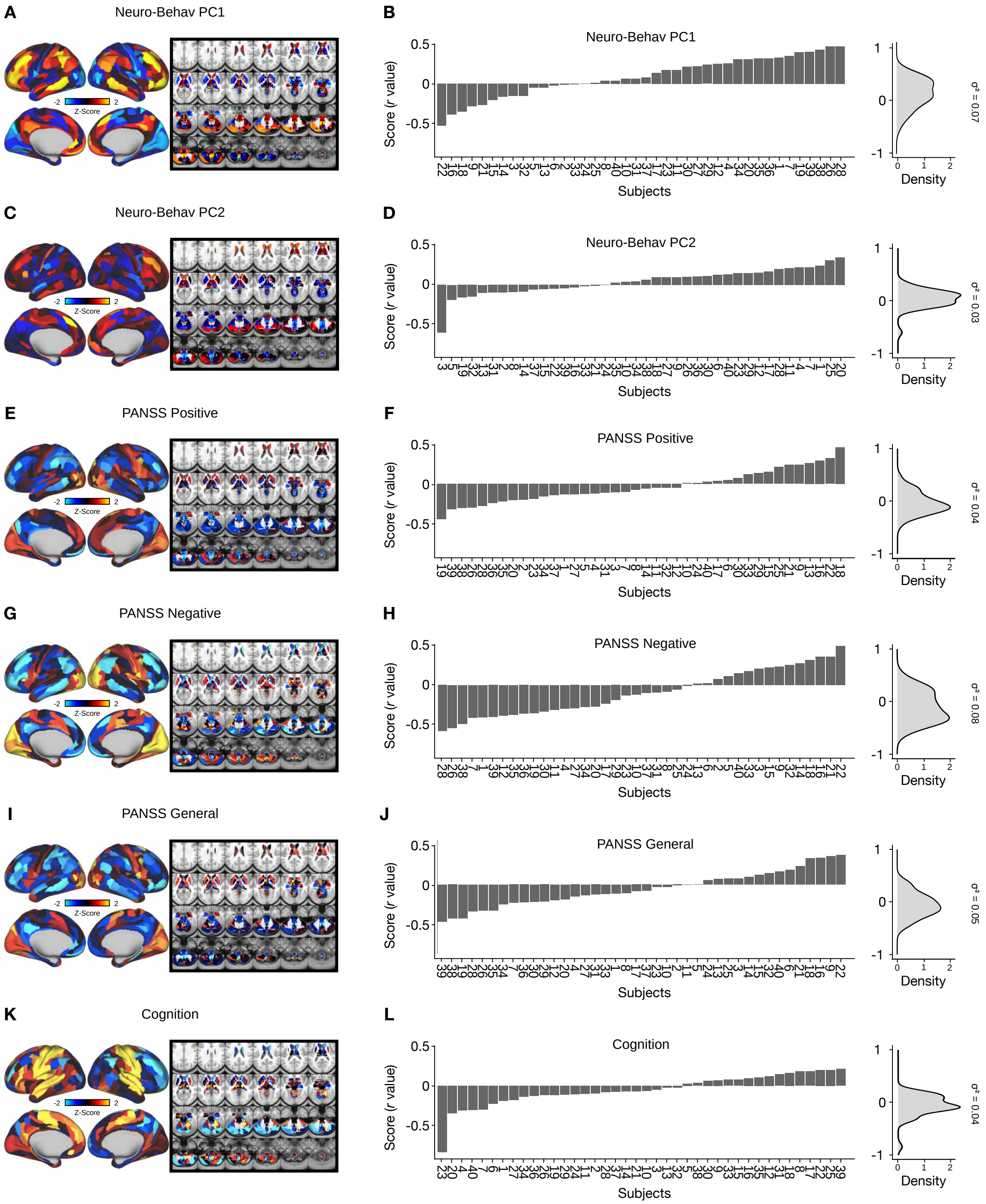

**Fig. S14. Individual variation captured by neuro-behavioral PCs, 3-factor PANSS subscales, and cognition Δ GBC maps. (A)** Neuro-behavioral PC1 map showing the relationship between the behavioral PC1 score for each participant regressed onto the Δ GBC map for each participant (N=40). **(B)** (Left) Bar plot showing the neuro-behavioral PC1 score (r) for each participant (N=40). Neuro-behavioral PC1 scores were calculated by correlating each participant's individual Δ GBC map with the neuro-behavioral PC1 Δ GBC map. (Right) density plot of neuro-behavioral PC1 scores. **(C)** Neuro-behavioral PC2 map showing the relationship between the behavioral PC2 score for each participant regressed onto the Δ GBC map for each participant. **(D)** (Left) Bar plot showing the neuro-behavioral PC2 score (r) for each participant (N=40). Neuro-behavioral PC2 scores were calculated by correlating each participant's individual Δ GBC map with the neuro-behavioral PC2 Δ GBC map. (Right) density plot of neuro-behavioral PC2 scores. **(E)** PANSS positive map showing the relationship between the PANSS positive score for each participant regressed onto the Δ GBC map for each participant. **(F)** (Left) Bar plot showing the PANSS positive score (r) for each participant (N=40). PANSS positive scores were calculated by correlating each participant's individual Δ GBC map with the PANSS positive Δ GBC map. (Right) density plot of PANSS positive scores. **(G)** PANSS negative map showing the relationship between the PANSS negative score for each participant regressed onto the Δ GBC map for each participant. **(H)** (Left) Bar plot showing the PANSS negative score (r) for each participant (N=40). PANSS negative scores were calculated by correlating each participant's individual Δ GBC map with the PANSS negative Δ GBC map. (Right) density plot of PANSS negative scores. **(I)** PANSS general map showing the relationship between the PANSS general score for each participant regressed onto the Δ GBC map for each participant. **(J)** (Left) Bar plot showing the PANSS general score (r) for each participant (N=40). PANSS general scores were calculated by correlating each participant's individual Δ GBC map with the PANSS general Δ GBC map. (Right) density plot of PANSS general scores. **(K)** Cognition map showing the relationship between the cognition score for each participant regressed onto the Δ GBC map for each participant. **(L)** (Left) Bar plot showing the cognition score (r) for each participant (N=40). Cognition scores were calculated by correlating each participant's individual Δ GBC map with the cognition Δ GBC map. (Right) density plot of cognition scores.

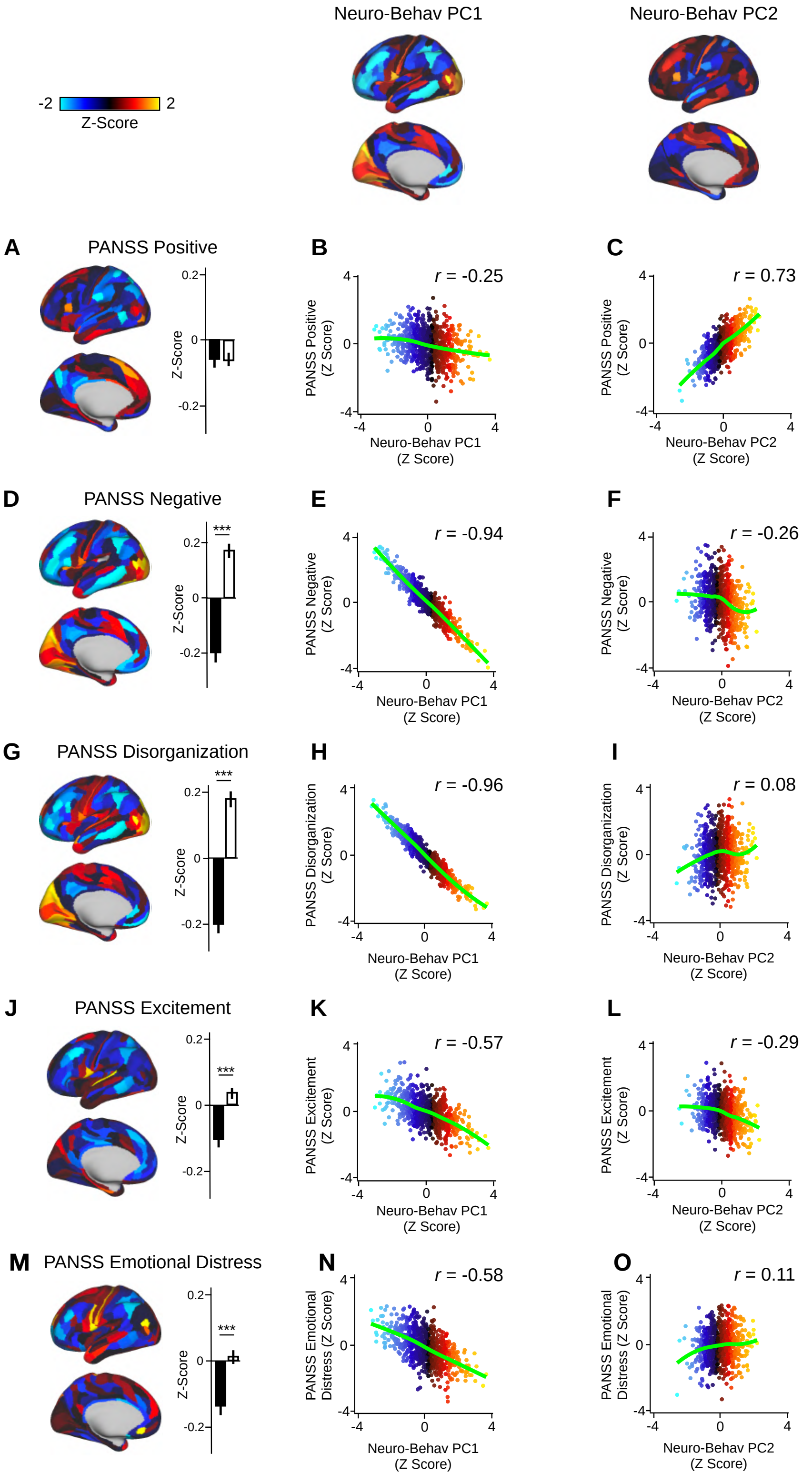


**Fig. S15. Scatter plots showing the relationship between neuro-behavioral PC1-2 and the 5-factor PANSS subscales. (A, D, G, J, M)** (left) Maps showing the relationship between the 5-factor PANSS subscale score for each participant regressed onto the △ GBC map for each participant (N=40). Values shown in each brain parcel are the Z-scored regression coefficient (PANSS subscale, △ GBC) across all 40 subjects. Red/orange areas indicate parcels in which there is a positive relationship between GBC and the PANSS subscale score, while blue areas indicate parcels in which there is a negative relationship between GBC and the PANSS subscale score. (right) Bar plot showing the mean Z-scored correlation value (PANSS subscale, △ GBC) for association (black) and sensory (white) networks. **(B)** Scatter plot showing the relationship between PANSS positive (left) and neuro-behavioral PC1 (top). **(C)** Scatter plot showing the relationship between PANSS positive (left) and neuro-behavioral PC2 (top). **(E)** Scatter plot showing the relationship between PANSS negative (left) and neuro-behavioral PC1 (top). **(F)** Scatter plot showing the relationship between PANSS negative (left) and neuro-behavioral PC2 (top). **(H)** Scatter plot showing the relationship between PANSS disorganization (left) and neuro-behavioral PC1 (top). **(I)** Scatter plot showing the relationship between PANSS disorganization (left) and neuro-behavioral PC2 (top). **(K)** Scatter plot showing the relationship between PANSS excitement (left) and neuro-behavioral PC1 (top). **(L)** Scatter plot showing the relationship between PANSS excitement (left) and neuro-behavioral PC2 (top). **(N)** Scatter plot showing the relationship between PANSS emotional distress (left) and neuro-behavioral PC1 (top). **(O)** Scatter plot showing the relationship between PANSS emotional distress (left) and neuro-behavioral PC2 (top).

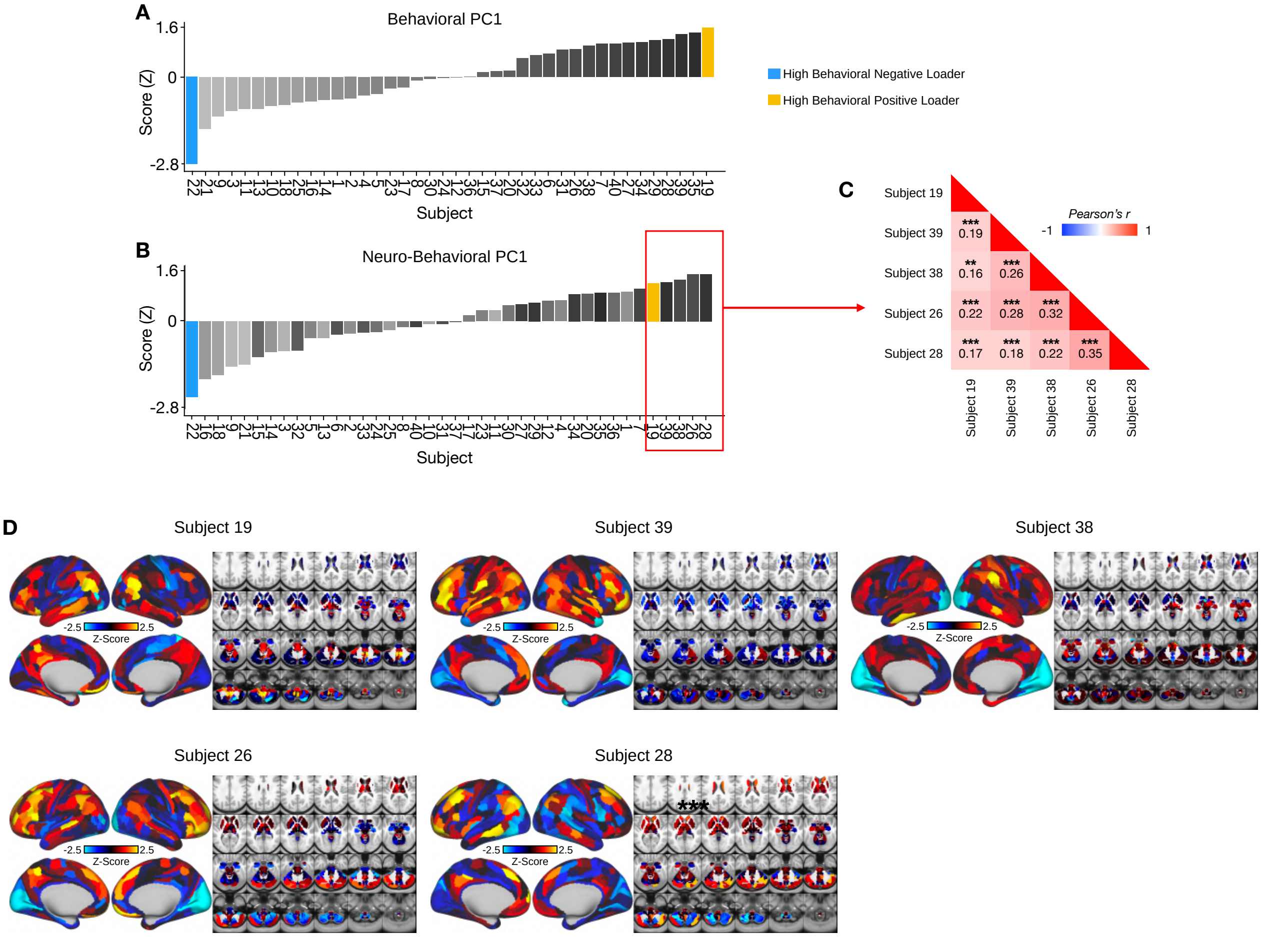


**Fig. S16. Comparison of the top five positive neuro-behavioral PC1 loaders. (A)** Bar plot showing the behavioral PC1 score (Z) for each individual participant (N=40). Bars are ordered and color-coded according to each participant's behavioral PC1 score (light grey = highly negative score, dark gray = highly positive score). Blue = high negative behavioral PC1 loader, yellow = high positive behavioral PC1 loader. **(B)** Bar plot showing the neuro-behavioral PC1 score (Z) for each individual participant (N=40). The neuro-behavioral PC1 score is calculated by correlating each participant's $\Delta$ GBC map with the neuro-behavioral PC1 map, and then z-scoring the r-values. Bars are ordered according to each participant's neuro-behavioral PC1 score, but color-coded according to each participant's behavioral PC1 score (light grey = highly negative score, dark gray = highly positive score). Blue = high negative behavioral PC1 loader, yellow = high positive behavioral PC1 loader. Top 5 positive loaders are highlighted in red. **(C)** Correlation matrix showing the relationship between the top five neuro-behavioral PC1 positive loaders' individual $\Delta$ GBC maps. *** = p<.001, ** = p<.01. **(D)** Top five neuro-behavioral PC1 positive loaders' individual $\Delta$ GBC maps. Red/orange areas indicate regions where participants exhibited stronger GBC in the ketamine condition, whereas blue areas indicate regions where participants exhibited reduced GBC in the ketamine condition.

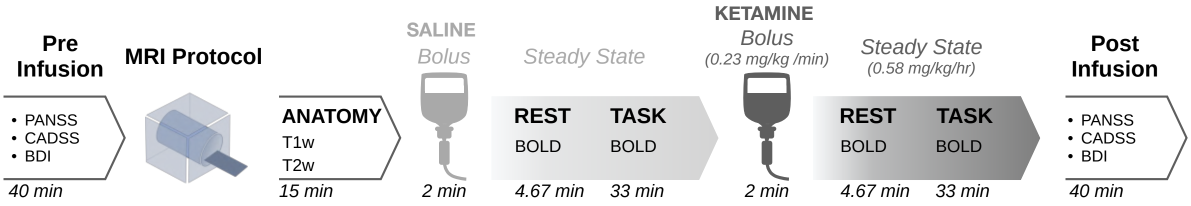


**Fig. S17. Neuroimaging Protocol.** The study employed a double-blind within-subjects design. 40 participants received an IV cannulation in each forearm: one IV for the placebo/ketamine infusion and one for blood draws during the scan. Placebo was administered during the first neuroimaging can session, and ketamine (initial bolus 0.23 mg/kg, continuous infusion 0.58 mg/kg/hour) during the second because residual ketamine effects ruled out counterbalancing of the order. Blood was drawn immediately after the resting-state run. Cognitive effects were measured using a Spatial Working Memory task completed in the scanner. Subjective effects were measured 180 mins post drug administration using the following scales: (1) Positive and Negative Syndrome Scale (PANSS), (2) Clinician Administered Dissociative States Scale (CADSS), and (3) Beck's Depression Inventory (BDI).

DRAFT

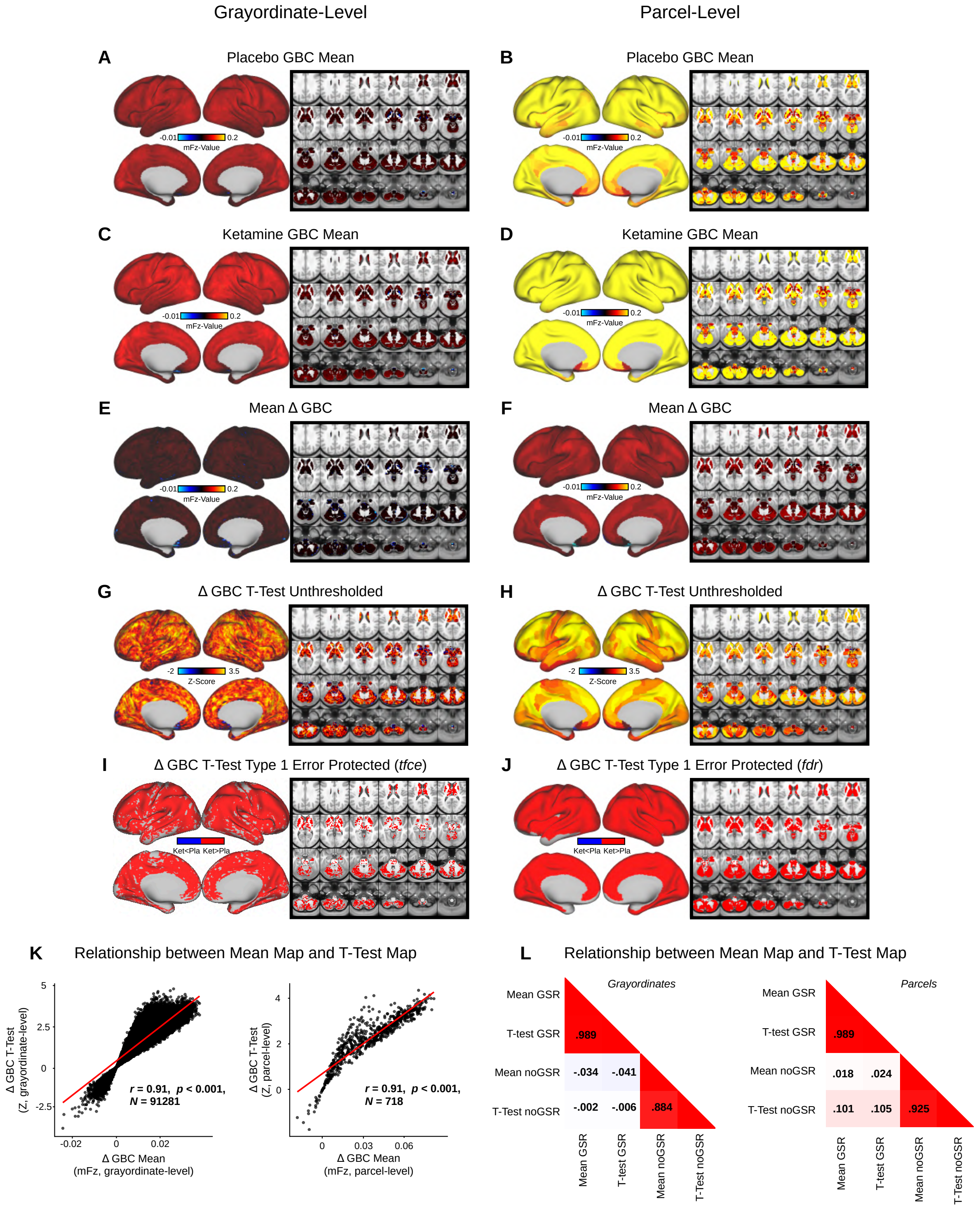


**Fig. S18. Mean effect of ketamine on global brain connectivity without global signal regression.** **(A-B)** Placebo mean GBC neural mFz-value map at the grayordinate level (left, no. grayordinates = 91281) and parcel level (right). **(C-D)** Ketamine mean GBC neural mFz-value map at the grayordinate level (left) and parcel level (right). **(E-F)** Mean Δ (ketamine - placebo) GBC neural mFz-value map at the grayordinate level (left) and parcel level (right). **(G-H)** Ketamine vs. placebo t-test unthresholded Z-score map at the grayordinate level (left) and parcel level (right). Red/orange areas indicate regions where participants exhibited stronger GBC in the ketamine condition, whereas blue areas indicate regions where participants exhibited reduced GBC in the ketamine condition. **(I-J)** Ketamine vs. placebo t-test significant (FDR protected, 5000 permutations) areas showing increased (red) and decreased (blue) GBC in the ketamine condition compared to placebo at the grayordinate level (left) and parcel level (right). **(K)** Scatterplots showing the relationship between the mean Δ GBC mFz-value map and the unthresholded t-test Δ GBC Z-score map (Z-scores computed across 718 parcels) at the grayordinate level (left) and parcel level (right). **(L)** Correlation matrix showing the relationship between mean Δ GBC map and t-test Δ GBC map with and without GSR at the grayordinate level (left) and parcel level (right).

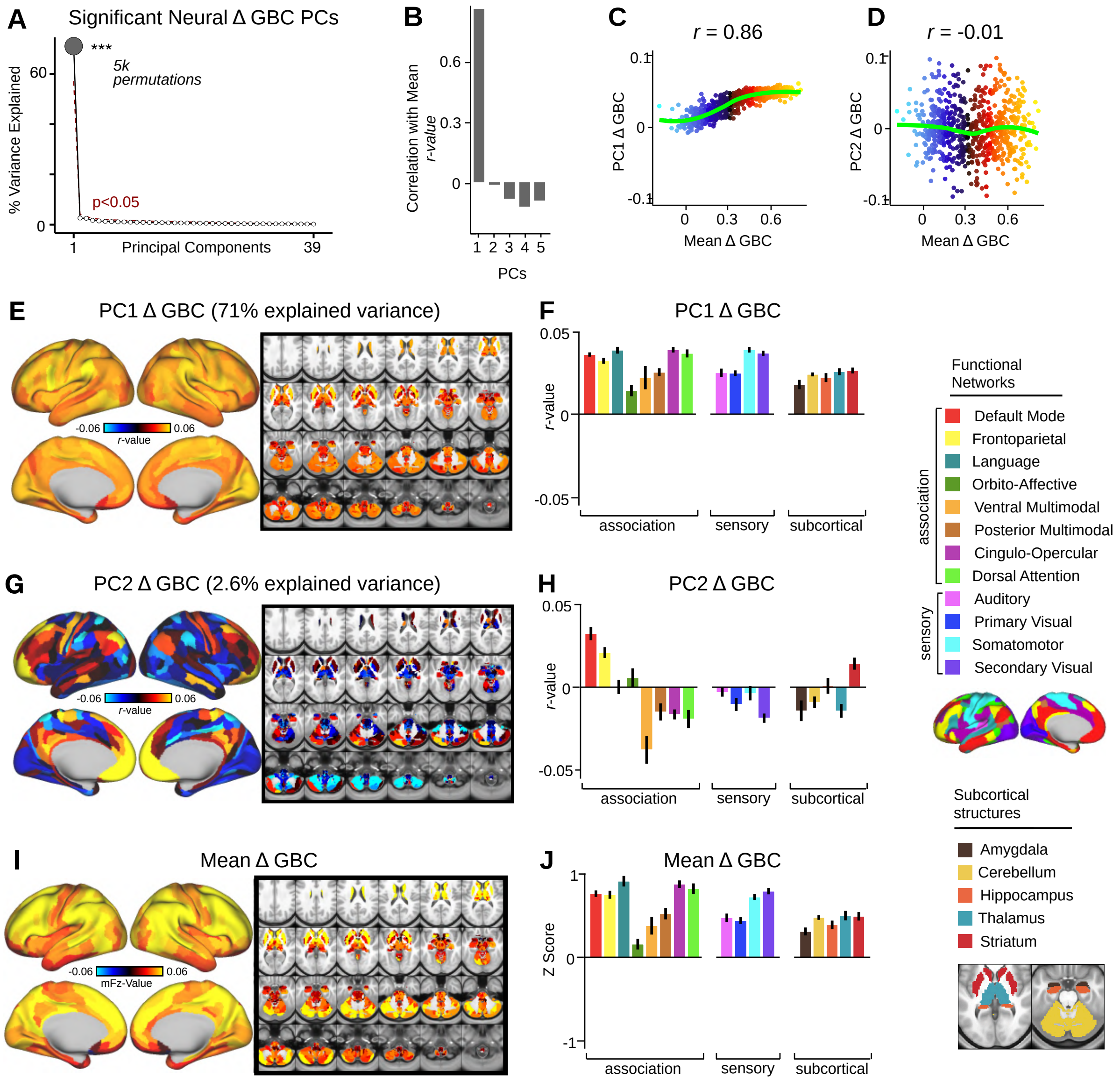


**Fig. S19. Multi-dimensional neural effect of acute ketamine administration without global signal regression. (A)** Results of PCA performed on Δ GBC neural features without global signal regression (GSR) (718 whole-brain parcel GBC) across all subjects (N=40). Screeplot showing the % variance explained by the 39 Δ GBC PCs. The first Δ GBC PC (dark grey) was determined to be significant using a permutation test (p<.05, 5000 permutations). PC1 Δ GBC captures 71% of the total variance in neural GBC in the sample. **(B)** Bar plot showing the correlation between the first five Δ GBC PCs and the mean Δ GBC. PC1 Δ GBC is most highly correlated with mean Δ GBC, which is to be expected as PC1 Δ GBC explains the largest amount of variance in the neural data and is the only significant PC. **(C)** Scatterplot showing the relationship across parcels between mean Δ GBC and PC1 Δ GBC maps (r=0.86, p<.001, N=718). Green line indicates that while the positive values are correlated, the negative values do not correlate. **(D)** Scatterplot showing the relationship across parcels between mean Δ GBC and PC2 Δ GBC maps (r=-0.01, p>.05, N=718). Green line indicates neither the positive or negative values are highly correlated. **(E)** PC1 Δ GBC r-value map. PC1 Δ GBC explains 71% of all variance. Red/orange areas indicate parcels that have a high positive loading score onto PC1, while blue areas indicate parcels that have a high negative loading score onto PC1. **(F)** Bar plot showing the mean correlation (Δ GBC, PC1 score) for each network and each anatomical subcortical structure for PC1 Δ GBC (see inset right for color labels). **(G)** PC2 Δ GBC r-value map. PC2 Δ GBC explains 2.6% of all variance. Red/orange areas indicate parcels that have a high positive loading score onto PC2, while blue areas indicate parcels that have a high negative loading score onto PC2. **(H)** Bar plot showing the mean correlation (Δ GBC, PC2 score) for each network and each anatomical subcortical structure for PC2 Δ GBC (see inset right for color labels). **(I)** Unthresholded mean Δ GBC mFz-value map. Red/orange areas indicate regions where participants exhibited stronger GBC in the ketamine condition, whereas blue areas indicate regions where participants exhibited reduced GBC in the ketamine condition, compared with the placebo condition. **(J)** Bar plot showing the mean Δ GBC mFz-value for each network and each anatomical subcortical structure for mean Δ GBC (see inset right for color labels).

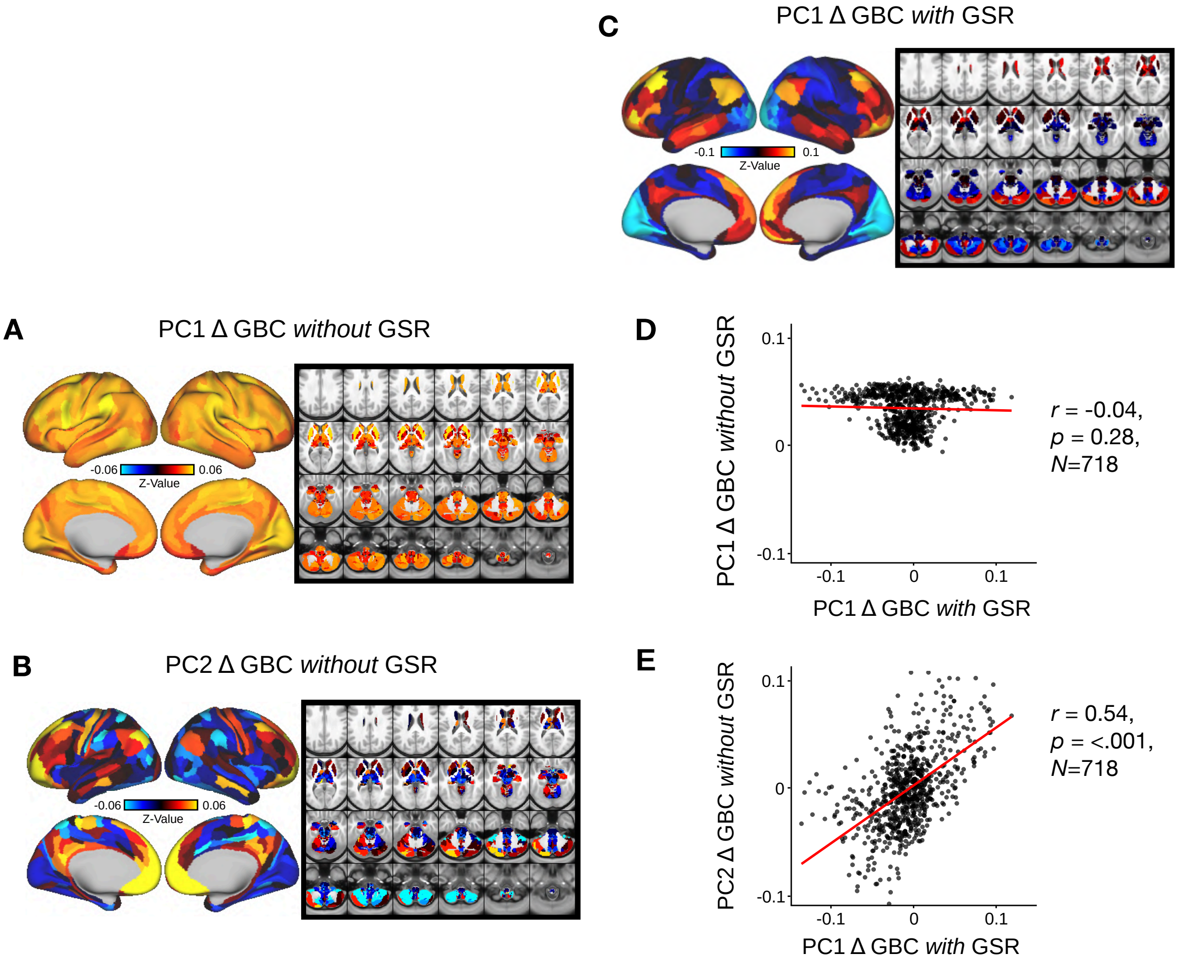


**Fig. S20. Comparison of Delta GBC Principal Component Analysis (PCA) with vs. without Global Signal Regression (GSR). (A)** PC1 △ GBC z-value map without GSR. PC1 △ GBC without GSR explains 71% of all variance. Red/orange areas indicate parcels that have a high positive loading score onto PC1, while blue areas indicate parcels that have a high negative loading score onto PC1. **(B)** PC2 △ GBC z-value map without GSR. PC2 △ GBC without GSR explains 2.6% of all variance. Red/orange areas indicate parcels that have a high positive loading score onto PC2, while blue areas indicate parcels that have a high negative loading score onto PC2. **(C)** PC1 △ GBC Z-score map with GSR (Z-scores computed across 718 parcels). PC1 △ GBC with GSR explains 14.1% of all variance. Red/orange areas indicate parcels that have a high positive loading score onto PC1, while blue areas indicate parcels that have a high negative loading score onto PC1. (PC1 is sign flipped for visual comparison with mean). **(D)** Scatter plot showing the correlation across parcels (N=718) between PC1 △ GBC Z-score map with GSR and PC1 △ GBC Z-score map without GSR (Z-scores computed across 718 parcels). **(E)** Scatter plot showing the correlation across parcels (N=718) between PC1 △ GBC Z-score map with GSR and PC2 △ GBC Z-score map without GSR (Z-scores computed across 718 parcels).

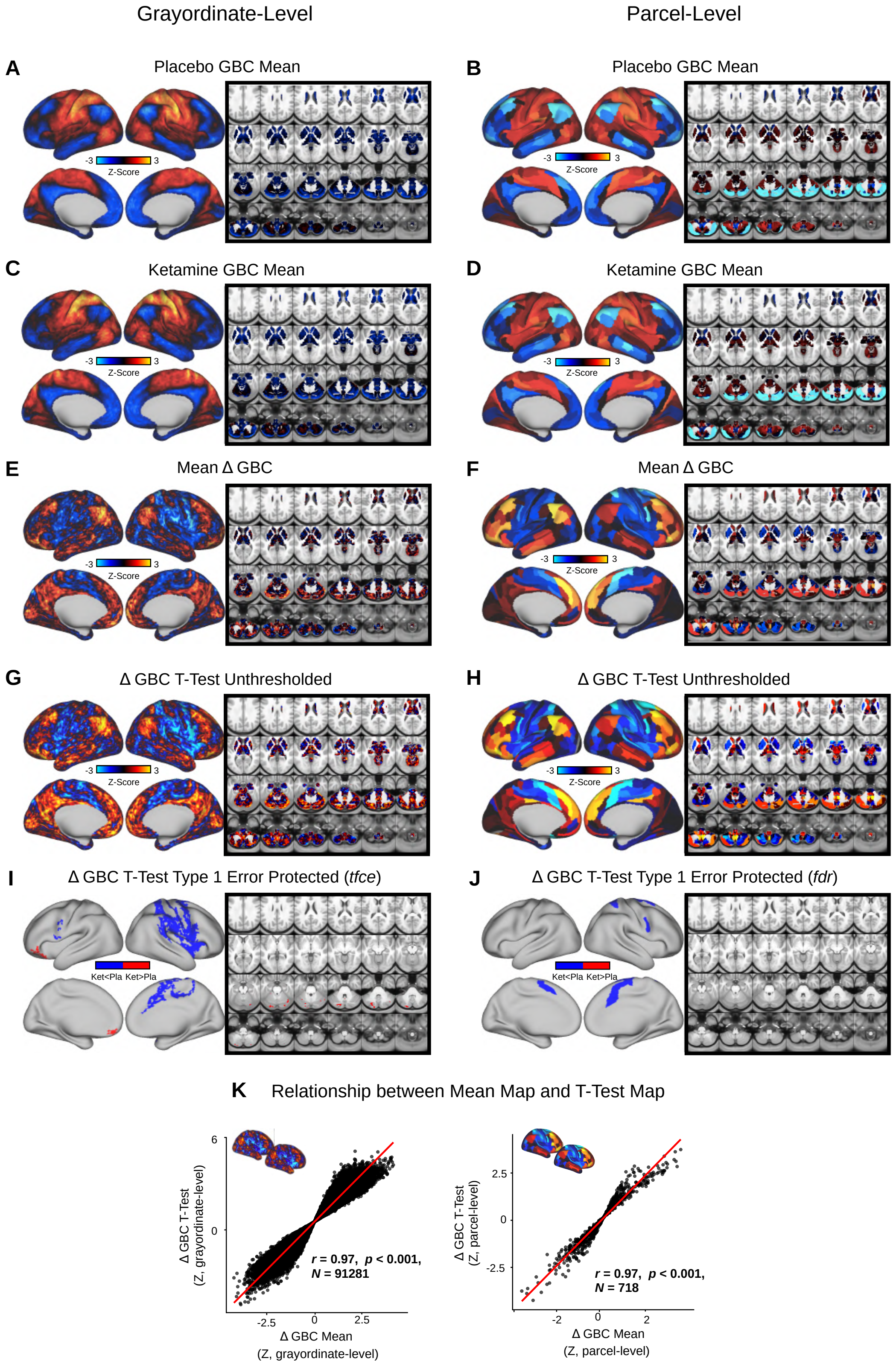


**Fig. S21. Mean effect of ketamine on GBC. (A-B)** Placebo mean GBC neural map at the grayordinate level (left, no. grayordinates = 91281) and parcel level (right). **(C-D)** Ketamine mean GBC neural map at the grayordinate level (left) and parcel level (right). **(E-F)** Mean Δ (ketamine - placebo) GBC neural map at the grayordinate level (left) and parcel level (right). **(G-H)** Ketamine vs. placebo t-test unthresholded Z-score map at the grayordinate level (left) and parcel level (right). Red/orange areas indicate regions where participants exhibited stronger GBC in the ketamine condition, whereas blue areas indicate regions where participants exhibited reduced GBC in the ketamine condition. **(I-J)** Ketamine vs. placebo t-test significant (threshold free cluster enhancement type 1 error protected, 5000 permutations) areas showing increased (red) and decreased (blue) GBC in the ketamine condition compared to placebo at the grayordinate level (left) and parcel level (right). **(K)** Scatterplots showing the relationship between the mean Δ GBC map and the unthresholded t-test Δ GBC map at the grayordinate level (left) and parcel level (right).
